# A Cross-tissue Temporal Multi-omics Atlas Reveals the Molecular Architecture of Pressure Injury

**DOI:** 10.64898/2026.05.21.727040

**Authors:** Fa Jiang, Liuyang Li, Yutong Zhao, Ting Zhang, Xue Li, Junchao Wei, Xiaona Liu, Yaru Jia, Meiwen An, Xiong Jiao

## Abstract

Pressure injury induces progressive necrosis of skin and deep muscle, yet how mechanical loading reshapes molecular programs across tissues and time remains poorly defined. To address this gap, we established a rat pressure-injury model and performed longitudinal integrated profiling of skin and deep muscle across four stages, combining transcriptomics, proteomics and untargeted metabolomics with histological assessment. This multi-layered atlas revealed a staged injury program dominated by early transcriptional activation of innate immunity and complement-coagulation crosstalk, accompanied by neutrophil-associated responses and engagement of upstream regulatory networks. As injury advanced, a second axis of remodeling emerged, characterized by sustained suppression of mitochondrial energy metabolism and oxidative phosphorylation, particularly in muscle, together with extracellular matrix and adhesion rewiring. In parallel, skin preferentially activated barrier-repair and protein-homeostasis programs, including keratinization and autophagy-linked processes, indicating tissue-specific adaptation to the same mechanical insult. Temporal clustering and pathway-network analysis further showed that immune activation and metabolic collapse are not independent events but are dynamically coupled across stages, with muscle exhibiting broader molecular reprogramming and a stronger shift toward irreversible structural failure. Together, these data define a cross-tissue, stage-resolved molecular framework for pressure injury progression and identify pathway-level windows that may inform tissue-targeted and stage-specific intervention strategies.

## Introduction

Pressure injury (PI), also known as pressure ulcer, is a clinical condition characterized by localized damage to the skin and underlying soft tissues (including subcutaneous fat and muscle) caused by sustained or repeated compression and shear forces. It commonly arises over bony prominences such as the sacrum, calcaneus and ischial tuberosity, and may present as intact skin with underlying tissue damage or as open ulcers. PI frequently affects individuals with limited mobility or sensory impairment, including hospitalized older adults, people with spinal cord injury and patients in intensive care [1, 2]. Epidemiological data indicate a global prevalence of approximately 12.8% for pressure injuries among hospitalized adults [3], and a risk ratio of 0.31 for hospital-acquired pressure injuries (HAPIs) [4]. In the United States alone, the annual direct medical cost attributable to PIs has been estimated at approximately US$17.8 billion [2]. Beyond the substantial physiological and psychological burden for affected patients, PI imposes major demands on healthcare resources consumption and constitutes a significant economic burden. In 2016, the National Pressure Ulcer Advisory Panel (NPUAP) updated the staging system for pressure injury, defining categories that include stages 1–4, unstageable injury, deep tissue injury and medical device- or mucosa-related injury, thereby improving clinical recognition, nursing quality assessment and comparability in research [5].

Under mechanical stress conditions, in addition to directly causing damage to cellular and tissue structures, it can also disrupt relevant signaling pathways and molecular expression, thereby indirectly participating in the initiation, progression, and repair of PI. Currently, four principal pathophysiological hypotheses have been proposed to explain the formation of PI, including local ischemia, reperfusion injury, impaired lymphatic drainage, and direct cellular mechanical deformation. However, these mechanisms are not isolated from each other, but interact across different tissue depths and temporal scales, ultimately precipitating sustained inflammatory responses, cellular necrosis, and impaired repair [6–8]. The precise molecular events and regulatory networks underlying these processes remain poorly understood. With the development of high-throughput sequencing and mass spectrometry technologies, multi-omics approaches (transcriptomics, proteomics, metabolomics, etc.) offer novel tools for systematically resolve the molecular dynamics of PI [9]. However, existing omics studies predominantly focus on single time points or isolated ‘layers’ [10, 11]. Furthermore, due to the difficulty in obtaining early clinical samples, individual heterogeneity, and differences in study designs, comparability between studies and the feasibility of translating findings to clinical practice remain challenging [12]. Consequently, establishing animal models that replicate the temporal progression of human disease and enabling longitudinal sampling across multiple molecular levels are crucial for elucidating key molecular events and regulatory networks during disease progression, thereby guiding stage-specific intervention strategies.

To comprehensively elucidate the molecular mechanisms underlying the development of pressure injuries, this study established and validated an animal model in rats encompassing four critical injury stages (with model reproducibility and staging characteristics confirmed through macroscopic observation and histopathology). Building upon this model, we conducted a longitudinal, integrated multi-omics analysis across multiple temporal scales, combining transcriptomics, proteomics and untargeted metabolomics with systematic data quality control and integrative analyses to construct a temporal molecular atlas of injury progression. Compared with prior single-time-point or single-omics studies, our work substantially enhances temporal resolution and tissue breadth. In conclusion, this study employs a multi-level, longitudinal systems biology approach to systematically depict the molecular evolution landscape of PI across different tissues and disease stages, providing actionable molecular clues and resources for subsequent targeted, stage-specific intervention strategies.

## Materials and Methods

### Pressure-injury animal model

Twenty-eight male Sprague Dawley (SD) rats (8 weeks old, weighing approximately 250 g) were randomly assigned to four groups (control s0, experimental s1, s2, s3), with seven rats per group. After a 1-week acclimation period, hair on the right hindlimb was shaved and the area disinfected with 75% ethanol. For experimental groups, two magnets (each 10 mm in diameter, 4 mm thick, with a magnetic flux density of 0.39 T) were placed on either side of the gracilis muscle of the right hindlimb. Each compression cycle consisted of 2 h of compression followed by 2 h of release, repeated three cycles per day. Animals in s1, s2 and s3 were euthanized and sampled at day 3, day 7 and day 14 after the start of the experiment, respectively; control animals (s0) were sampled at the end of the acclimation period and served as baseline controls. Tissue sampling was performed immediately after euthanasia: wound tissue was dissected, skin and underlying muscle separated and allocated as follows: a portion rinsed in ice-cold PBS, snap-frozen in liquid nitrogen and stored at −80°C for sequencing; a portion fixed in 4% paraformaldehyde for histology; remaining material retained as reserve. Animals were singly housed with ad libitum access to food and water; ambient temperature was maintained at 25 ± 2°C and relative humidity at 40–50%. Body weight was recorded at the experimental start and at each animal’s endpoint. Multi-group body weight comparisons were performed by one-way ANOVA with Tukey HSD post-hoc test; within-group baseline versus endpoint comparisons used paired t-tests. Significance was set at two-sided p < 0.05. This study has been approved by the ethics committee. All animals were supplied by the Laboratory Animal Center of Shanxi Medical University (License No.: SCXK (Jin) 2024-0004). The study protocol was approved by the Biomedical Ethics Committee of Taiyuan University of Technology (Approval No.: TYUT2025122303).

### Histopathology

Fixed tissues were processed by standard dehydration, paraffin embedding and sectioning (4 µm). Sections were floated on water at 40°C, mounted and baked at 60°C until dry. Sections were stained with hematoxylin–eosin (H&E) and Masson’s trichrome by standard procedures and examined by light microscopy; images were captured for analysis. Image analysis was performed in Fiji [13] (ImageJ). Details of histopathological quantification are provided in the Supplementary Methods.

### Sequencing and data processing

Transcriptome. Sequencing was performed by Novogene. Raw reads were processed to remove adapter sequences, poly-N sequences and low-quality reads to obtain high-quality clean data. Clean reads were aligned to the Ensembl Rattus norvegicus (mRatBN7.2) reference genome using HISAT2 (v2.2.1). Gene counts were obtained with featureCounts (v2.0.6). Low-expressed genes were removed (counts per million < 0.5 in all but one sample). The filtered count matrix was analyzed using DESeq2 [14] and normalized with the variance-stabilizing transformation (vst). TMM normalization was performed using calcNormFactors from the package edgeR [15] and converted to counts per million (CPM) for transcript-level coefficient of variation (CV) calculations.

Proteome. Sequencing was performed by Novogene. Raw data were searched with DIA-NN against the UniProt database. Search parameters included automatic correction of precursor and fragment ion mass errors, fixed modification: cysteine alkylation; variable modification: N-terminal methionine oxidation; up to two missed cleavages. Filtering thresholds were global peptide Q-value < 0.01 and protein Q-value < 0.01. Protein abundance was normalized by total sum normalization. Proteins with > 1/3 missing values across samples were excluded; remaining missing values were imputed using K-nearest neighbors (KNN) method from the R package impute. Quantitative data were log2-transformed for downstream analysis.

Metabolome. Sequencing was performed by Novogene. Raw files were converted to mzXML (ProteoWizard) and processed by XCMS software for peak extraction, alignment and quantitation (mass tolerance 10 ppm). Metabolite identification was based on secondary mass spectrometry database matching. Quantification was normalized using quality control (QC) samples as follows: Normalized value = Raw quantitative value of sample / (Sum of quantitative values of metabolites in sample / Sum of quantitative values of metabolites in QC1 sample). Features with CV > 30% in QC samples were removed. Final data were log2-transformed for downstream analysis. Further procedural detail is in the Supplementary Methods.

### Outlier detection and dimensionality reduction

Principal component analysis (PCA) was performed separately on each omics layer (DESeq2 vst for transcriptome; log2-normalized data for proteome and metabolome) to determine variance explained by each principal component (PC). Outliers were defined as samples with scores on any PC explaining > 7.5% of variance that lay outside [Q1 − 3×IQR, Q3 + 3×IQR], where Q1 and Q3 are the first and third quartiles and IQR is the interquartile range. For dimensionality reduction, PCA was applied to z-score standardized data (stats::prcomp); for joint multi-omics analyses, t-distributed stochastic neighbor embedding (t-SNE) was applied to the z-score standardized merged data (Rtsne package) with perplexity = 5, theta = 0.05 and a fixed random seed.

### Molecular variability

CV was calculated on non-imputed data for each molecular feature as CV = (Standard deviation / Mean) × 100%. For transcriptomics CVs were computed using CPM; for proteomics and metabolomics CVs were computed on pre-imputation, non-log standardized data. Definitions of high-variability features were: (1) Stage-specific high variability: CV > 100% at that stage; (2) Post-injury high variability: median CV across s1–s3 > 100%; (3) Relative baseline variability: (Post-injury CV / Baseline s0 CV) > 300%. Technical variability in metabolomics was assessed using QC samples.

### ID mapping and cross-species mapping

Transcript Ensembl IDs were mapped to Gene Symbols and Entrez IDs; proteome UniProt IDs were mapped to Gene Symbols and Entrez IDs; metabolome HMDB IDs were mapped to KEGG IDs. Mapping used the rat genome annotation file (https://download.rgd.mcw.edu/data_release/RAT/GENES_RAT.txt, accessed 1/6/2025) to relate Gene Symbols, Ensembl IDs, Entrez IDs and RGD IDs.

Rat-human orthologue mapping used the RGD orthologue file (https://download.rgd.mcw.edu/data_release/RAT/ORTHOLOGS_RAT.txt, accessed: 16/9/2025) to relate rat Ensembl IDs, Entrez IDs, Gene Symbols, RGD IDs to human gene identifiers.

### Differential expression analysis

Differential analyses were performed independently for each omics layer and tissue. For transcriptomics, inter-group comparisons used DESeq2 on filtered gene counts; the model included group as the main effect and RNA integrity number (RIN) as a covariate. For proteomics and metabolomics, log2-transformed normalized data were analyzed with limma [16] using empirical Bayes variance shrinkage. To screen for molecules altered throughout the entire injury process, transcriptomics used the likelihood ratio test (DESeq2::nbinomLRT), comparing a full model (group + RIN) to a reduced model (RIN only); proteomics and metabolomics used an F-test in limma. P-values were adjusted by Benjamini–Hochberg false discovery rate (FDR), with significance defined as FDR < 0.05.

### Enrichment analyses

Pathway gene sets were integrated from KEGG [17] and Reactome [18] databases after removing redundant pathways (≥80% feature overlap). Background sets comprised all detected molecules in each omics layer. Transcript and protein IDs were mapped to Entrez IDs; metabolites to KEGG IDs. Enrichment analysis used the R package clusterProfiler [19], incorporating the aforementioned custom background molecular sets. Unless otherwise specified, pathway enrichment used the deduplicated combined pathway set (metabolite enrichment employed KEGG pathways only). Enrichment significance was set at FDR < 0.05. For molecular variability analysis, pathways significantly enriched in highly variable molecules at the transcript or protein level were required to contain more than five molecules. For enrichment analysis of molecular interaction network clusters within each trajectory, significant pathways were required to contain more than one molecule.

Gene set enrichment analysis (GSEA) was performed using the GSEA function within the clusterProfiler R package. Input data comprised the log2FC-ordered list derived from differential analysis. Gene sets included the aforementioned deduplicated KEGG and Reactome collections and a custom HMDB metabolite classification set. Enrichment significance was set at FDR < 0.05.

Differentially expressed genes (FDR < 0.05) from each tissue were subjected to transcription factor motif enrichment with HOMER [20] (v5.1) using findMotifs.pl in Ubuntu 20.04.6. The enrichment level of sequence motifs within tissues was quantified by the percentage of differentially expressed gene promoters containing the motif. Enrichment p-values were calculated by comparing enrichment levels in the target genome against a background genome normalized for GC content and corrected by the Benjamini method to obtain q-values; significance threshold q-value < 0.05. Hierarchical clustering based on the mean absolute difference value of TF enrichment differences across tissues assessed similarity in motif enrichment patterns between tissues.

Rat feature IDs (transcripts and proteins) were mapped to human Entrez IDs via the rat–human orthologue mapping. Differential features (FDR < 0.05) were subjected to Disease Ontology enrichment using DOSE::enrichDO [21]. Terms with FDR < 0.05 were considered significantly enriched.

### Bayesian feature clustering

Bayesian soft clustering and trajectory construction were performed on the z-scores or t-scores (hereafter denoted as z) from the differential expression analysis results, utilizing the R package repfdr [22]. Let *Z* ∈ *R^n×t^*denote the input score matrix, where *z_i,j_* represents the z-value for feature *i* within injury group *j* ∈ {1, …, t} (in this study, t=3 corresponds to groups s1, s2, s3). Assume each observation *z_i,j_* originates from a mixture of two distributions: null (no difference, *H* = 0, *N*(0,1)) and non-null (difference exists, *H* ∈{−1, 1} denoting downregulation or upregulation respectively). Introduce a latent state *h_i,j_* ∈ {-1,0,1} for each position; The complete state vector h for each feature *i* is a vector of length t, representing the directional pattern of that feature across all groups. The repfdr Expectation-Maximization (EM) algorithm is run as follows: First, for all z-values collected across features in each group *j*, repfdr::ztobins was used for binning and estimating null and non-null densities, with a manually selected degree of freedom of 20;

Subsequently, repfdr::repfdr is employed for EM, jointly estimating the prior probability p(*h*) of all possible complete states *h* and the posterior probability Pr(*h*|*z_i_*) of each feature in each state. To mitigate noise effects, the set of states with prior probability p(*h*) below 0.001 is discarded. Groups enumeration yields s1, s2, s3, with s0 added as a starting node containing all features to facilitate tracing the origin. Define node S*(j*,*o*) as the set of group *j* in state *o* ∈ {-1, 0, 1}. Calculate the probability of feature *i* belonging to group S(*j*,*o*) by summing the posterior probability Pr(*h*|*z_i_*) for all complete states *h* where the *j*th feature equals *o*. If this sum exceeds the threshold θ = 0.5, feature *i* is assigned to node S(*j*,*o*). Construct a set of edges between adjacent time points to represent the temporal evolution of features. For nodes in adjacent groups *j* and *j*+1, directly take the intersection of the node sets as the edge set.

### Pathway enrichment networks and community construction

For trajectory-specific features, pathway enrichment was performed as described above (FDR < 0.05). For pathways enriched in multiple groups, p-values were combined using the Fisher’s sum of logs method. Similarity between pathways was calculated based on the scoring criteria of Cytoscape [23] Enrichment Map, defined as follows: Jaccard = [size of (A intersect B)] / [size of (A union B)], Overlap = [size of (A intersect B)] / [size of (minimum (A, B))], Similarity score = 0.5 × Jaccard + 0.5 × Overlap. Pathways were considered functionally similar and connected only when the similarity score ≥ 0.5. Community clustering was performed on each pathway using igraph::cluster_louvain, based on modularity metrics and hierarchical methods. The functional annotation of community networks was performed manually according to the types of pathways contained within each community.

### Molecular interaction network

Gene–gene and gene–metabolite interactions were extracted from KEGG and Reactome (rat) using KEGGREST and Reactome resources, without considering interaction directionality. Metabolite IDs from different sources were mapped to RefMet IDs via Metabolomics Workbench [24] (https://www.metabolomicsworkbench.org/); gene IDs from different sources were mapped to RGD IDs. The rat KEGG molecular network comprised 8,088 nodes and 76,240 edges; the Reactome comprised 5,430 nodes and 21,435 edges. Following network merging and deduplication, the rat molecular background network contained 10,209 nodes and 96,675 edges. Specific subgraphs were generated from the background network based on features within each trajectory. Transcript and protein features were collapsed to common gene nodes, for gene–gene edges we retained node pairs whose median Spearman correlation across all samples was ≥ 0.5. Module partitioning used igraph::cluster_leading_eigen (leading eigenvector algorithm); modules with more than 10 nodes were visualized in Cytoscape and pathway enrichment analysis was conducted for molecules within each module.

### Immune cell deconvolution

Immune composition in bulk transcriptomes was estimated with CIBERSORTx [25] (v1.05; https://cibersortx.stanford.edu/) using the LM22 signature matrix (22 immune cell types). The transcriptome count matrix was normalized to transcripts per million (TPM) and rat Ensembl IDs converted to human Gene Symbols via the orthologue file prior to running CIBERSORTx. Parameters: B-mode batch correction, quantile normalization disabled, absolute mode enabled, and 1000 permutations. Absolute scores were employed for inter-group statistical comparisons; relative scores (absolute score / sum of absolute scores per sample) visualized the relative proportions of immune cells within each sample. Expression levels of key immune cell marker genes at the transcript or protein level were used for auxiliary validation.

### Similarity to human disease

To evaluate the translational relevance of this animal model, published human pressure injury (PI) data were incorporated for systematic cross-species comparison. Histological images were obtained from Richards et al. [26] (human PI patient skin, H&E staining, from Supplementary Fig. 1A of the cited publication) and Liu et al. [10] (human PI patient muscle, H&E staining, from Fig. 1A of the cited publication) for morphological comparison. Proteomic comparisons were based on proteome sequencing data from PI patients and normal muscles reported by Liu et al. [27]: human protein identifiers were converted to rat gene identifiers via the rat–human orthologue mapping file and only successfully mapped features were retained. Subsequently, Spearman’s correlation coefficients between log2FC values for corresponding features across both species were calculated to assess overall trend consistency. Overlap significance of differential features was evaluated using Fisher’s exact test (two-sided); directional concordance among overlapping differentially features was tested by an exact binomial test (two-sided). All significance tests employed a two-tailed p < 0.05 threshold; differential features were defined using a strict threshold of FDR < 0.05 and |log2FC| > 1.

**Fig. 1.**
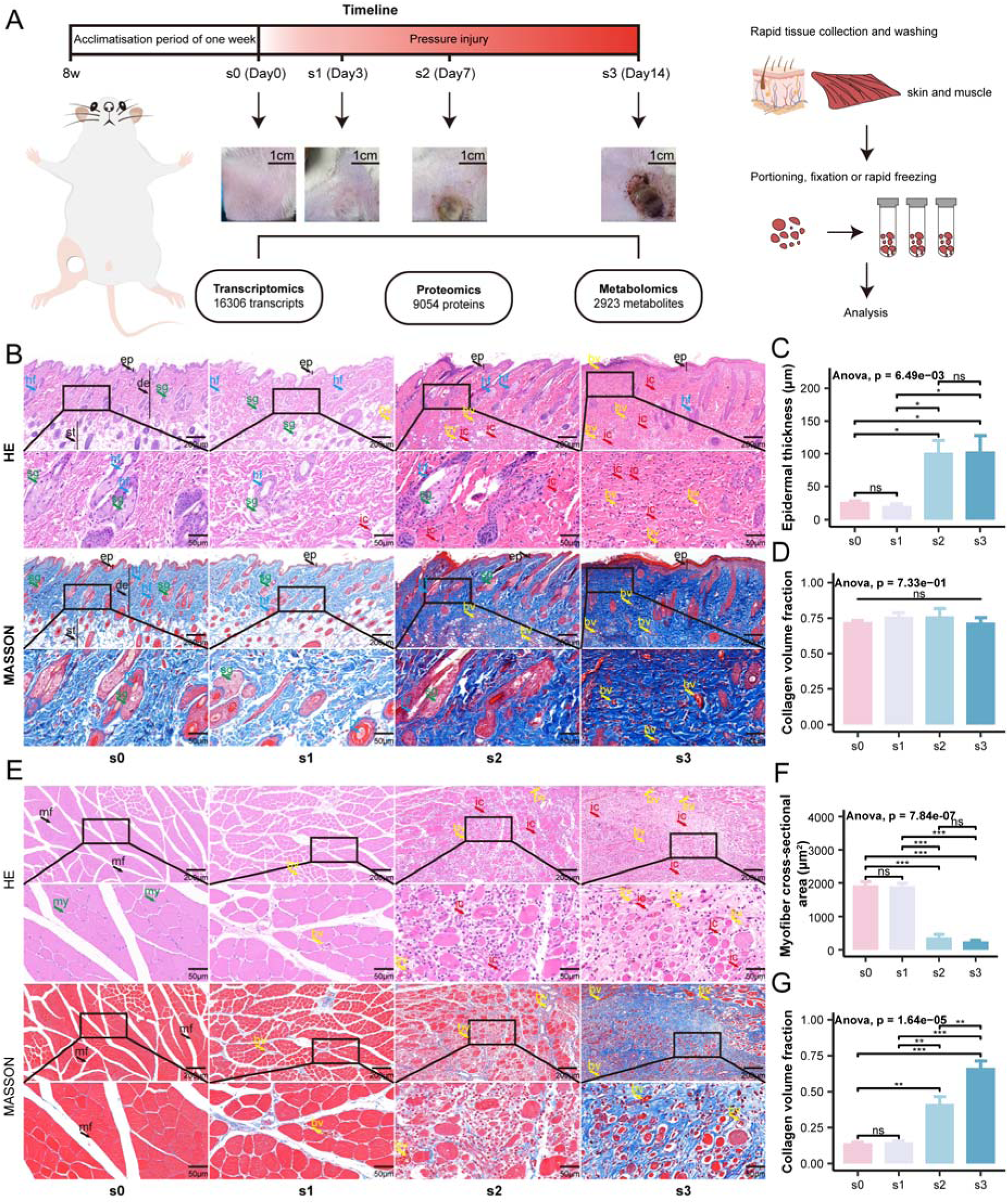
Study design and animal model assessment. (**A)** Schematic of the experimental workflow. Eight-week-old male Sprague–Dawley rats were acclimated for one week and then subjected to cyclic compression–release for 0 (s0), 3 (s1), 7 (s2) or 14 (s3) days. Tissues were collected at each terminal timepoint, partitioned and stored for downstream sequencing, histology and biobanking. (**B)** Representative skin H&E and Masson images for each group (top: scale bar, 200 µm; bottom: scale bar, 50 µm). (**C**) Epidermal thickness in skin for each group (n = 3). (**D)** Collagen volume fraction (CVF) in skin for each group (n = 3). (**E)** Representative muscle H&E and Masson images for each group (top: scale bar, 200 µm; bottom: scale bar, 50 µm). Abbreviations in **B**, **E**: ep, epidermis; de, dermis; st, subcutaneous tissue; hf, hair follicle; sg, sebaceous gland; bv, blood vessel; ic, inflammatory cell; mf, muscle fiber; my, myonucleus. (**F)** Muscle fiber cross-sectional area (CSA) for each group (n = 3). (**G)** Muscle collagen volume fraction (CVF) for each group (n = 3). Data in **C**, **D**, **F** and **G** are presented as mean ± SEM. Significance markers: ns, p ≥ 0.05; *p < 0.05; **p < 0.01; ***p < 0.001.

### Statistical analysis

All statistical analyses were performed using R software (v4.3.1). Specific sample sizes (n), statistical test methods, data description formats (mean ± standard deviation SD or standard error SEM), and definitions of significance levels are detailed in the corresponding methods section, main text or figure legends.

## Results

### Assessment of the pressure injury animal model

Following one week of acclimatization in 8-week-old Sprague–Dawley rats, a pressure injury (PI) model was established by applying varying numbers of compression–release cycles to the gracilis muscle of the right hindlimb. Macroscopic observations revealed: normal tissue appearance at baseline s0 (Day 0); localized skin discoloration, hyperemia, and pronounced indentation without skin breakdown at s1 (Day 3); full-thickness skin loss with exposure of subcutaneous fat, without involvement of the underlying muscle at s2 (Day 7); and full-thickness skin loss with muscle exposure, extensive necrotic tissue and eschar formation at s3 (Day 14) (Fig. 1A). These macroscopic features corresponded to normal tissue, Stage1 (intact skin with non-blanching erythema), Stage 3 (full-thickness skin loss with visible fat), and Stage 4 (full-thickness skin and tissue loss with exposed fascia, muscle, etc.) in human clinical staging [5]. For clarity, timepoints 0, 3, 7 and 14 days are hereafter designated s0, s1, s2 and s3, respectively.

Between-group comparisons of body weight at the start and endpoint revealed no significant differences; however, paired within-group comparisons showed a significant increase in body weight from start to endpoint for each group (paired t-test, p < 0.05; Supplementary Fig. 1A; Supplementary Table 1). Histopathology (Fig. 1B, E) revealed that in the s0 group, skin appendages and all layers exhibited intact structures with no evident inflammatory cell infiltration, and muscle fibers were regularly arranged with a homogeneous cross-sectional area. As the injury progressed, the epidermal layer structure remained discernible in the s1 stage, though the epidermal-dermal junction became lax. In s2-s3 stages, epidermal thickening and progressive destruction of skin appendages (hair follicles, sebaceous glands, etc.) were evident (Fig. 1C). Muscle tissue demonstrated disrupted myofibrillar structure, increased interstitial space, and rounded fiber morphology, accompanied by substantial inflammatory cell infiltration. Quantitatively, the tissue-stained area fraction (TAF) in skin increased at later injury stages, while the collagen volume fraction (CVF) showed no significant overall change, likely influenced by post-injury vascular hyperplasia and substantial inflammatory cell infiltration (Fig. 1D; Supplementary Fig. S1C). In muscle, the cross-sectional area of individual muscle fibers decreased during stages s2 and s3 concomitant with a marked increase in CVF, indicating progressive fibrosis, while TAF remained largely unchanged throughout the process (Fig. 1F, G; Supplementary Fig. S1D). Together, the above results demonstrate the validity of the pressure injury animal model established in this study and support subsequent multi-omics profiling. All omics datasets passed pre-specified quality-control filters (Supplementary Results).

### Molecular variability across different stages in individuals

To assess molecular heterogeneity across individuals and disease process, we calculated the coefficient of variation (CV) for molecules in each omics layer. In skin, variability increased post-injury relative to baseline for all omics except proteins; in muscle, variability increased post-injury across all omics. At baseline, CVs ranked from highest to lowest were: metabolism (negative ion mode), metabolism (positive ion mode), transcriptome, and proteome (approximately 35.9%, 30.0%, 27.3%, and 24.7% respectively); a similar ordering was observed post-injury (approximately 43.0%, 36.0%, 33.5%, and 23.5% respectively) (Fig. 2B; Supplementary Fig. 3A). In muscle, both metabolic positive and negative ion modes maintained high variability at baseline (approximately 39.6% and 41.1%) and post-injury (approximately 41.3% and 44.3%); transcriptome and proteome variability increased post-injury, with the transcriptome CV (approximately 34.0%) slightly exceeding that of the proteome (approximately 33.1%) (Fig. 2H; Supplementary Fig. 4A; Supplementary Table 3).

**Fig. 2.**
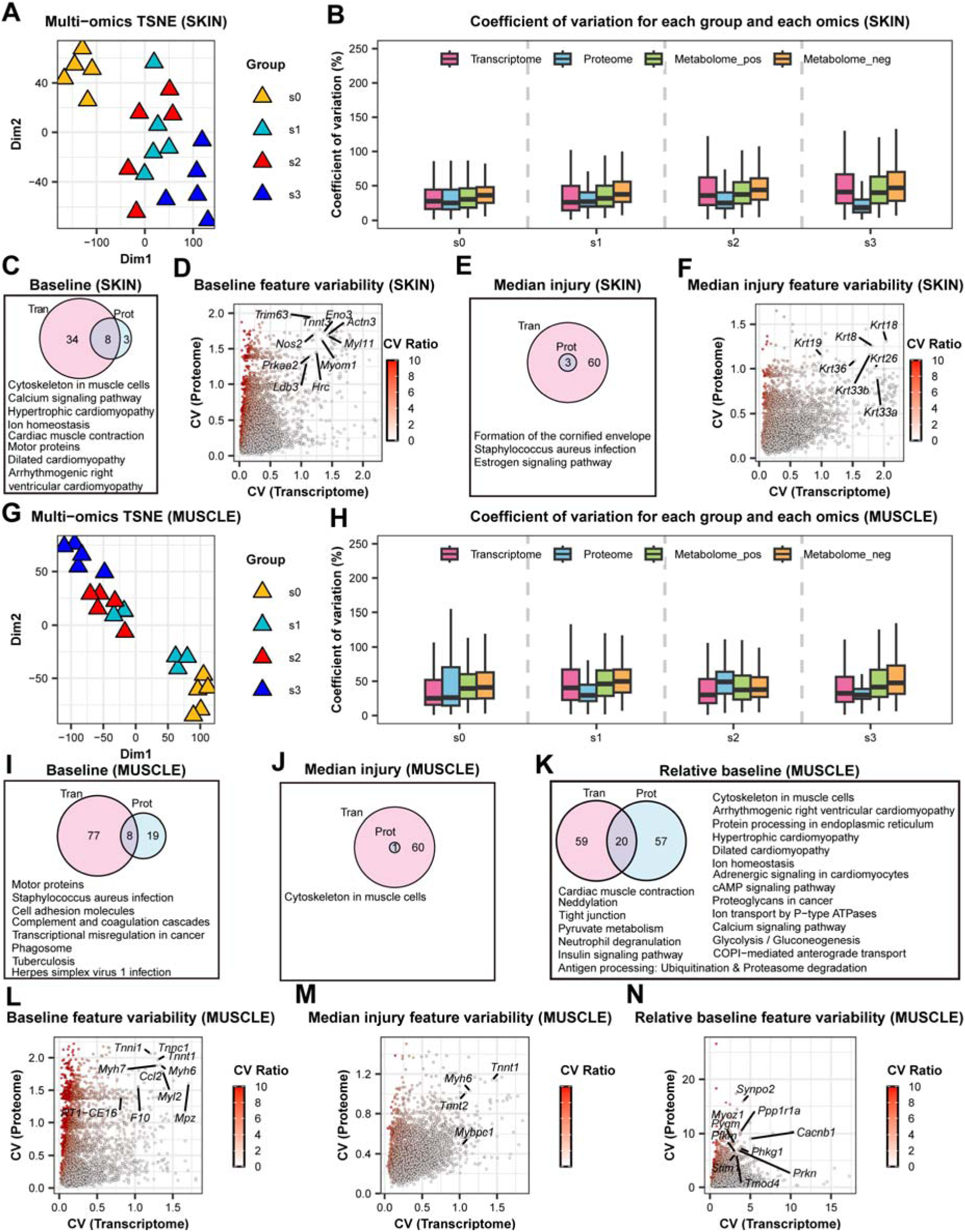
Molecular variability. (**A)** Two-dimensional t-distributed stochastic neighbor embedding (t-SNE) projection of skin samples based on combined multi-omics features; each point denotes one sample. (**B)** Boxplots of coefficient of variation (CV) for multi-omics features in skin across groups. (**C)** Venn diagram of pathways significantly enriched among high-CV transcripts and proteins at baseline (s0) in skin. (**D)** Comparison of CV between transcript and protein layers at baseline (s0) in skin; molecules that enter the same significant enrichment sets in both omics layers are highlighted and the CV ratio is shown as CV-protein / CV- transcript. (**E)** Venn diagram of pathways significantly enriched among high-CV transcripts and proteins in skin post-injury. (**F)** Post-injury transcript–protein CV ratio comparison in skin (annotation as in **D**). (**G–N)**, Corresponding analyses for muscle: **G**, muscle t-SNE; **H**, muscle CV boxplots; **I–K**, Venn diagrams of enriched pathways for high-CV transcripts and proteins at baseline, post-injury and relative-to-baseline conditions; **L–N**, transcript–protein CV ratio comparisons for the three conditions (annotation as in **D**). Boxplot notation: box limits define the first quartile Q1 (lower edge), median (center line), and third quartile Q3 (upper edge); whiskers denote non-outlier minimum and maximum.

We next calculated the proportion of highly variable molecules at each stage. In skin, transcriptomics contributed the largest proportion, followed by metabolomics and proteomics (approximately 16.9%, 15.1%, and 11.0%; Supplementary Fig. 3E). In muscle, metabolomics contributed most, followed by proteomics and transcriptomics (approximately 21.8%, 21.6%, and 19.6%; Supplementary Fig. 4E). Temporally, relative to baseline, skin exhibited greater transcriptional and proteomic fluctuations at s2, whereas metabolomics showed increased variability at s3. Muscle demonstrated most proteomic fluctuations at s2, with marked transcriptional and metabolomic changes at s3 (Supplementary Fig. 3F, 4F). The low CV values in metabolomics QC samples confirmed controlled technical variation (Supplementary Fig. 3B, 4B). Classification of metabolites detected in both positive and negative ion modes revealed that QC samples exhibited the lowest overall coefficient of variation, followed by baseline and post-injury samples (Supplementary Fig. 3C, D; Supplementary Fig.4C, D; Supplementary Table 4).

Functional enrichment analysis of highly variable molecules revealed that both transcriptomic and proteomic data at baseline were enriched in cytoskeletal pathways. Post-injury, both transcriptomic and proteomic layers were enriched for cornified envelope formation and keratin-related molecules (e.g., Krt18, Krt8) and exhibited high variability. These molecular alterations correlate with the histologically observed epidermal thickening, supporting the notion of injury- or inflammation-driven proliferative-differentiative reprogramming [28]. (Fig. 1C; Fig. 2E, F; Supplementary Fig. 3G, H). Relative to baseline, the C-type lectin receptor signaling pathway exhibited significant transcriptional fluctuations, and molecules within classical pathways such as JAK-STAT and TNF signaling also showed considerable variability. At the protein level, several metabolic processes (e.g., butyrate metabolism, fatty acid degradation) fluctuated (Supplementary Fig. 3I). Metabolomic fluctuations yielded fewer enriched pathways (Supplementary Fig. 3G-I). At baseline and post-injury, factors associated with receptor signaling exhibited significant transcriptional fluctuations in muscle tissue, accompanied by metabolic alterations; post-injury, cytoskeletal activity occurred at both the transcriptomic and proteomic levels within muscle cells, high variability was observed in myocyte-associated molecules such as myosin Myh6 and troponin Tnnt1, these molecules not only maintain normal muscle activity at baseline but also respond to injury stimuli. The observed cytoskeletal fluctuations, also present relative to baseline, may result from myofiber atrophy and fragmentation as seen in H&E and Masson staining. Relative to baseline, both transcriptional and protein levels of myocyte-associated disease molecules exhibited pronounced fluctuations. Additionally, molecular fluctuations in cAMP signaling, calcium signaling, and insulin signaling were observed (Fig. 1E; Fig. 2I-N; Supplementary Fig. 4G-I; Supplementary Table 5). Overall, injury-induced molecular fluctuations exhibit both shared features and marked tissue- and omics-specific differences.

### Multi-omics changes induced by pressure injury

Differential analyses revealed widespread molecular perturbations across all three omics layers. Inter-group differential analyses revealed that injury collectively induced significant alterations in 9,978 molecules within skin (approximately 36.1% of identified molecules, FDR < 0.05), comprising 5,262 transcripts (32.8%), 4,096 proteins (47.0%), and 620 metabolites (21.2%). Muscle exhibited greater changes with 17,059 significant molecules (approximately 69.0% of identified molecules, FDR < 0.05), comprising 10,526 transcripts (69.3%), 4,960 proteins (75.2%), and 1,573 metabolites (53.8%) Compared with baseline, skin had the most differentially expressed molecules at s2 (4,350 transcripts, 3,551 proteins, 426 metabolites), followed by s3 (2,296 transcripts, 1,761 proteins, 386 metabolites) and s1 (2,011 transcripts, 1,751 proteins, 210 metabolites); whereas muscle exhibited a progressive increase in differentially regulated molecules with injury severity (s1: 5628; s2: 12695; s3: 15396) (Fig. 3A, C).

**Fig. 3.**
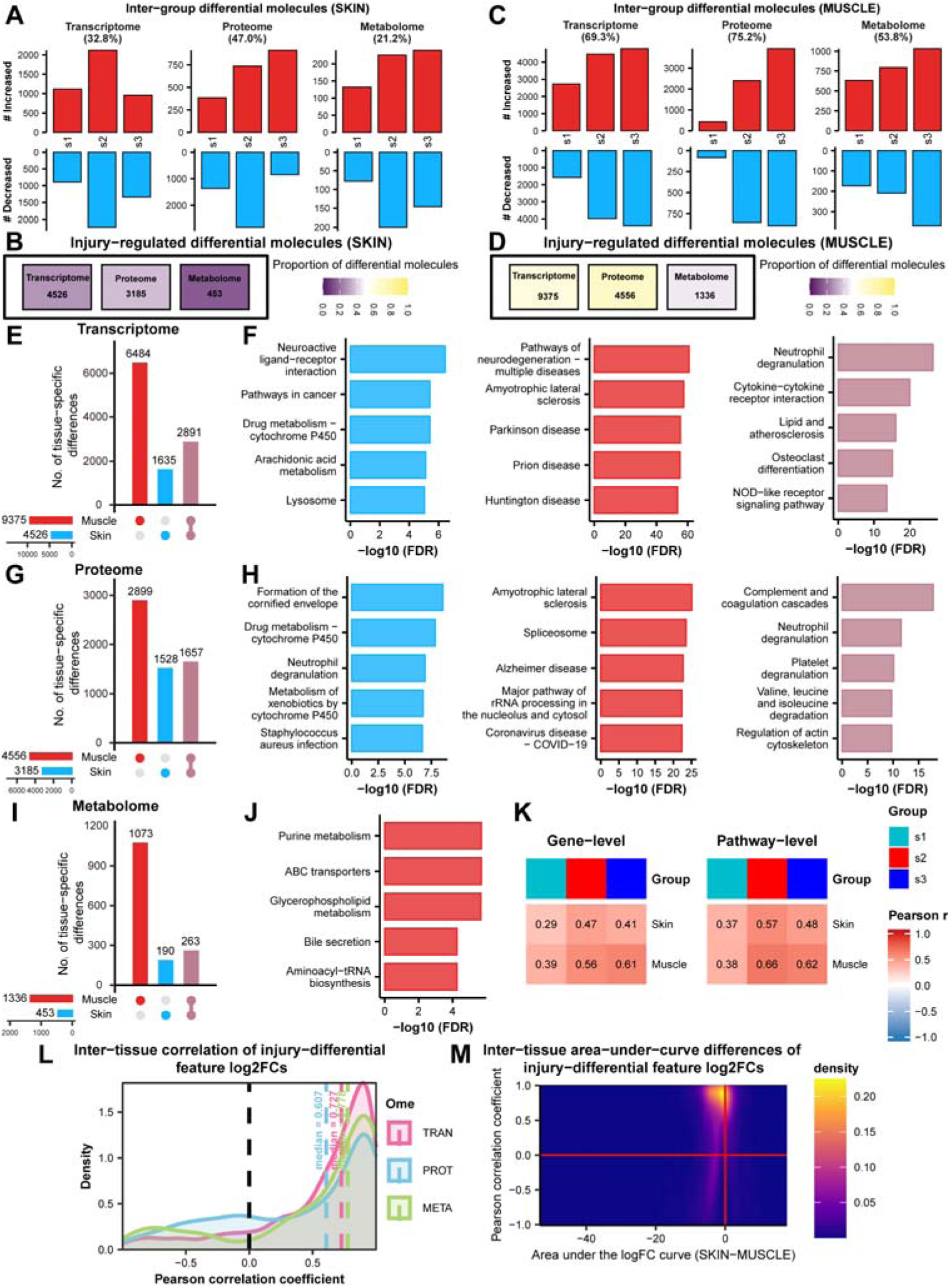
Shared and tissue-specific molecular and pathway changes between skin and muscle. (**A**) Counts of significant features (relative to s0) in skin for each injury stage (FDR < 0.05). (**B**) Total counts of significant features in skin across the full injury course (FDR < 0.05). (**C**) Counts of significant features (relative to s0) in muscle for each injury stage (FDR < 0.05). (**D**) Total counts of significant features in muscle across the full injury course (FDR < 0.05). (**E)** UpSet plot of transcripts with significant injury regulation in skin and muscle. Bars and connected dots denote tissue-specific or shared sets; colors indicate tissue membership. (**F)** Top five significantly enriched pathways for the transcript sets shown in **E** (left, skin-specific; center, muscle-specific; right, shared). (**G)** UpSet plot of significantly changed proteins (annotation as in **E**). (**H)** Top five significantly enriched pathways for the protein sets shown in **G** (annotation as in **F**). (**I)** UpSet plot of differential metabolites (annotation as in **E**). (**J)** Top five enriched metabolic pathway categories for muscle-specific differential metabolites (from **I**). (**K)** Heatmap of Pearson correlations between transcript-level and protein-level differential statistics (z-score and t-score respectively) and between normalized pathway enrichment scores (NES) from GSEA (left, gene level; right, pathway level). (**L)** Pearson correlation of log2FC values for shared differential features between tissues, shown separately for each omics layer. (**M)** Comparison of area under the curve (AUC) differences for log2FC curves of shared features between tissues. TRAN, transcriptome; PROT, proteome; META, metabolome.

Furthermore, statistical analysis of significantly altered molecules throughout the injury process revealed the proteome accounted for the highest proportion in both tissues (skin: 63.6%; muscle: 69.1%), followed by the transcriptome (skin: 28.2%; muscle: 61.7%) and the metabolome (skin: 15.5%; muscle: 45.7%). The transcriptome exhibited the highest number of altered molecules, followed by the proteome and finally the metabolome, a distribution consistent with the results of the inter-group differential analysis (Fig. 3B, D; Supplementary Table 6). Overall, muscle exhibited a larger magnitude of molecular reprogramming across omics in response to pressure injury.

### Tissue-specific molecular and pathway similarities and differences in pressure injury

Comparing differential molecules and pathway enrichments across tissues for the overall injury response revealed that muscle exhibited more tissue-specific changes across all three omics layers. Skin and muscle shared 4,811 altered molecules (2,891 transcripts; 1,657 proteins; 263 metabolites) (Fig. 3E, G, I). Transcriptome enrichment indicated skin-specific pathways included neuroactive ligand–receptor interaction, cytochrome P450–mediated drug metabolism, lysosome and IGF transport and regulation, whereas muscle-specific pathways included several neurodegenerative disease pathways, thermogenesis and oxidative phosphorylation; shared transcriptional enrichments included neutrophil degranulation, cytokine–cytokine receptor interaction and pathways such as NOD-like receptor and IL-17 signaling (Fig. 3F). Notably, neutrophil degranulation, an immune-regulatory signature, showed consistent transcriptional and proteomic upregulation and was persistently upregulated across the injury course by GSEA (Supplementary Fig. 5).

At the protein level, skin-specific enrichments centered on cornified envelope formation, cytochrome P450 metabolism and insulin signaling; muscle-specific enrichments involved muscular dystrophy-related pathways, rRNA processing and RNA splicing. Shared protein pathways included complement and coagulation cascades, neutrophil and platelet degranulation and actin cytoskeleton regulation (Fig. 3H). The most significantly enriched pathway, the complement and coagulation cascades (mediates innate immune recognition and hemostatic responses), exhibited a marked tendency towards high expression at the protein molecular level. GSEA results similarly demonstrated sustained upregulation of this pathway, suggesting its critical role in tissue injury and immune responses (Supplementary Fig. 6 and Supplementary Fig. 7C). At the metabolomic level, muscle displayed more tissue-specific enrichments (e.g., purine metabolism, ABC transporters, glycerophospholipid metabolism) (Fig. 3J; Supplementary Table 7).

Cross-tissue log2FC correlation analyses showed a positive correlation bias across omics (median correlations: transcriptome 0.727; proteome 0.607; metabolome 0.778). Furthermore, the area under the log2FC curve differences between tissues were predominantly concentrated near zero, indicating substantial overall concordance in molecular directionality between tissues (Fig. 3K-M). GSEA indicated sustained upregulation of many immune-related responses (chemokine signaling, complement, coagulation, neutrophil and leukocyte pathways) in both tissues, while metabolic pathways (e.g., mitochondrial protein degradation, fatty acid catabolism, pyruvate metabolism) were more markedly downregulated in muscle, suggesting impaired energy metabolism and potential hindrance to muscle repair (Supplementary Fig. 7C; Supplementary Table 8). These results illustrate the similarities and differences omics responses of the two tissues to pressure injury, underscoring the necessity and advantages of multi-tissue, multi-omics integration in deciphering the biological responses to pressure injury.

### Transcription factor motif enrichment analysis

We performed transcription factor (TF) motif enrichment analysis on differentially expressed genes to infer upstream regulators. Overall, TF motifs were significantly enriched primarily in the early injury stage (s1), suggesting an early regulatory influence on molecular changes. At s1, both tissues exhibited enrichment of ETS family motifs (e.g., PU.1 (also known as Spi1), Spib). Additionally, skin displayed Cebp-related motifs, which regulate gene expression involved in immune and inflammatory responses. These factors are associated with hematopoietic, immune cell function and inflammatory responses, particularly evident in B cells and dendritic cells. In contrast, muscle exhibited a broader repertoire of enriched TF families at s1 (including KLF and Fos families) linked to differentiation, stress responses and muscle regeneration. At s2, the significantly enriched TF Yy1 primarily governs cell differentiation and proliferation; Elk4 serves as a key cofactor for the serum response factor (SRF); Nfy functions in cell cycle progression, proliferation, and metabolic reprogramming; Sp2, a member of the SP family, plays a role in cell growth, differentiation, and apoptosis (Supplementary Fig. 8A; Supplementary Table 9).

Hierarchical clustering of transcription factor motif enrichment scores across different tissues at each stage revealed that enrichment profiles were relatively similar within tissues, whereas marked differences existed between tissues, reflecting tissue-specific transcriptional regulatory responses (Supplementary Fig. 8B). Further analysis integrating TF expression levels revealed that significantly differentially expressed factors exhibited an overall upregulation trend during the injury phase, clustering into distinct dynamic trajectories. Notably, the three encoded subunits constituting the Nfy complex (Nfya, Nfyb, Nfyc) displayed non-identical expression patterns: Nfya was significantly upregulated in injured groups, whereas Nfyb and Nfyc showed slight decreases at the same time point, suggesting potential modulation of Nfy function via subunit composition or relative abundance — that will require subsequent functional experiments to validate its actual impact on the complex’s DNA-binding activity and downstream transcriptional regulation (Supplementary Fig. 8C, D). Certain TFs exhibited a sharp early upregulation followed by a decline or return to baseline levels at s3 (e.g., Spi1 and Spib in skin; Fos and Klf4 in muscle), whereas others maintained high expression, indicating that the injury-induced transcriptional response comprised both transient immediate early responses and long-term transcriptional reprogramming.

### Molecular trajectories

To characterize the temporal dynamics of multi-omics molecules, we employed an empirical Bayesian approach for molecular clustering. The RepFDR diagnostic plot (three stages, degrees of freedom = 20) demonstrated that the empirical zero distribution effectively fitted the overall z-score distribution, whilst the quantile-quantile (Q-Q) plot exhibited significant deviation at both tails, indicating enrichment of genuine signals in the tails (Supplementary Fig. 9A, B). Molecules were ultimately grouped into ten temporal trajectories (Supplementary Table 10), with the first two trajectories containing the highest number of molecules across both tissues, corresponding to sustained upregulation and sustained downregulation (s1_1 -> s2_1 -> s3_1 and s1_-1 -> s2_-1 -> s3_-1). We focused on the top three trajectories per tissue, which together captured the majority of differential features (skin 79.3%: 6,463/8,153; muscle 87.5%: 13,360/15,263) to characterize principal dynamic patterns (Supplementary Fig. 9C, D).

In skin, the primary trajectory was sustained downregulation (SKIN: s1_-1 -> s2_-1 -> s3_-1), the secondary was sustained upregulation (SKIN: s1_1 -> s2_1 -> s3_1), and the third exhibited a decline followed by recovery to baseline at s3 (SKIN: s1_-1 -> s2_-1 -> s3_0). In muscle, the primary trajectory showed sustained upregulation (MUSCLE: s1_1 -> s2_1 -> s3_1), the second trajectory exhibited sustained downregulation (MUSCLE: s1_-1 -> s2_-1 -> s3_-1), and the third demonstrated initial stability followed by upregulation between s2 and s3 (MUSCLE: s1_0 -> s2_1 -> s3_1). The top three trajectories in skin contained a relatively balanced proportions of transcription and protein molecules, whereas in muscle the first two trajectories were dominated by transcript-level molecules and the third trajectory showing the highest proportion of protein molecules (Fig. 4).

**Fig. 4.**
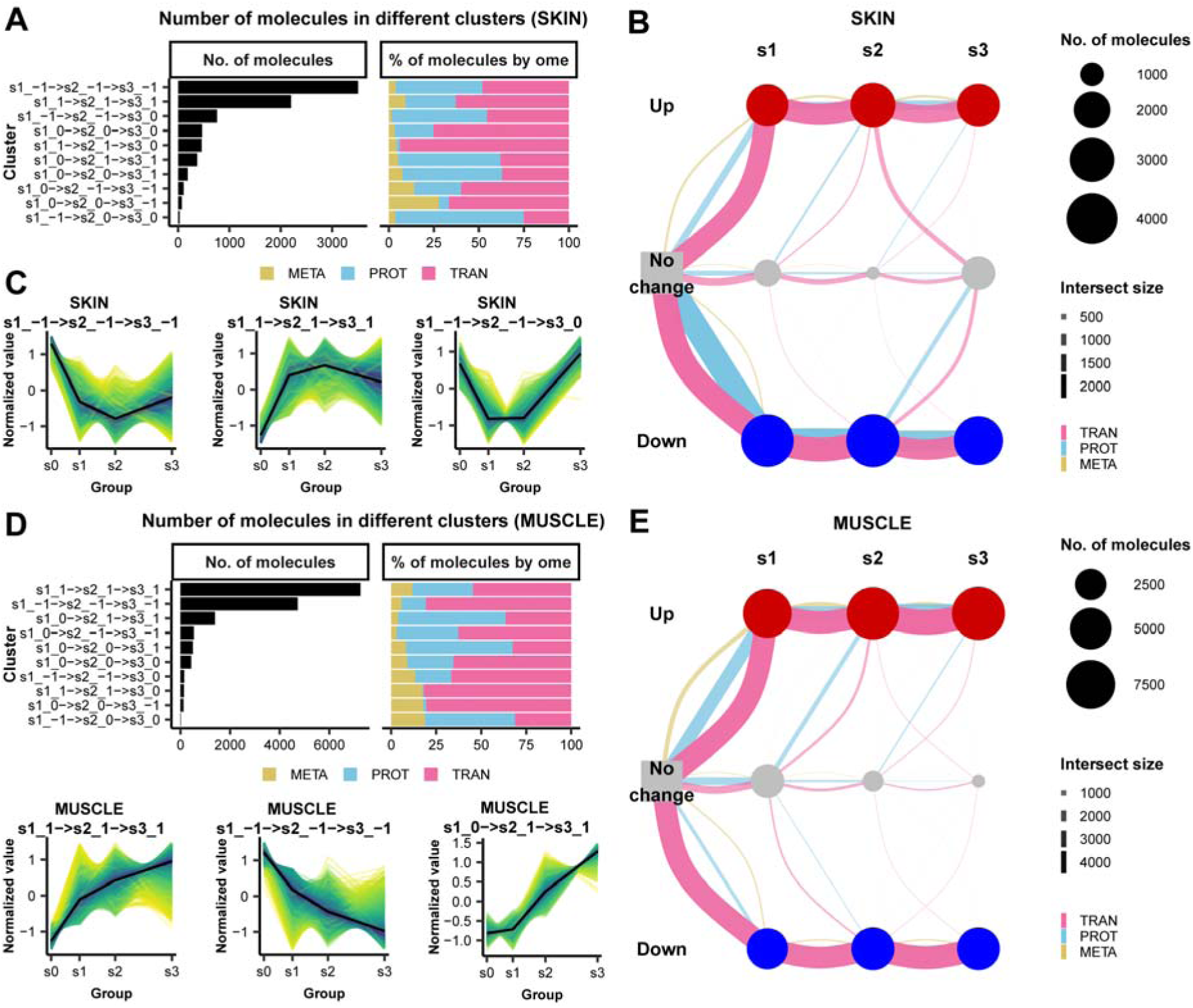
Temporal trajectory analysis of injury-regulated molecules. (**A)** Number of molecules assigned to each trajectory class in skin and the omics composition of each class. (**B)** Flow diagram for skin trajectories: nodes represent discrete stages and states; node size is proportional to the number of features assigned; edges represent transitions and edge width is proportional to the number of features following that trajectory. (**C)** Z-score-standardized expression profiles for the top three skin trajectories (those with the largest feature counts); the black line is the mean z-score across all features in the trajectory. (**D–F)** Corresponding analyses for muscle: **D** trajectory counts and omics composition; **E** trajectory flow diagram; **F** z-score profiles for the three largest muscle trajectories (black line denotes mean).

### Enriched pathways and interaction networks of molecules under different trajectories

#### Sustained up-regulated trajectory (s1_1 -> s2_1 -> s3_1)

Shared pathway enrichments in this trajectory concentrated on immune responses (e.g., complement and coagulation cascades, B cell-associated pathways), cell growth and death, RHO family signaling and carbohydrate metabolism-related pathways. Pathway-network analysis highlighted microtubule-C-terminal post-translational modification pathways as highly connected, suggesting roles for microtubule stability and transport in mechanical stress responses (Fig. 5A, B; Supplementary Table 12). In shared high-connectivity network nodes, Nfkb1 occupied a central position and co-enriched with chemokine and cytokine–receptor pathways (cluster 1). Other shared core nodes included Syk, Jak2, Shc1 and Vav1, implicating coordinated immune and signaling activation across clusters (Fig. 5C; Supplementary Fig. 13; Supplementary Table 13).

**Fig. 5.**
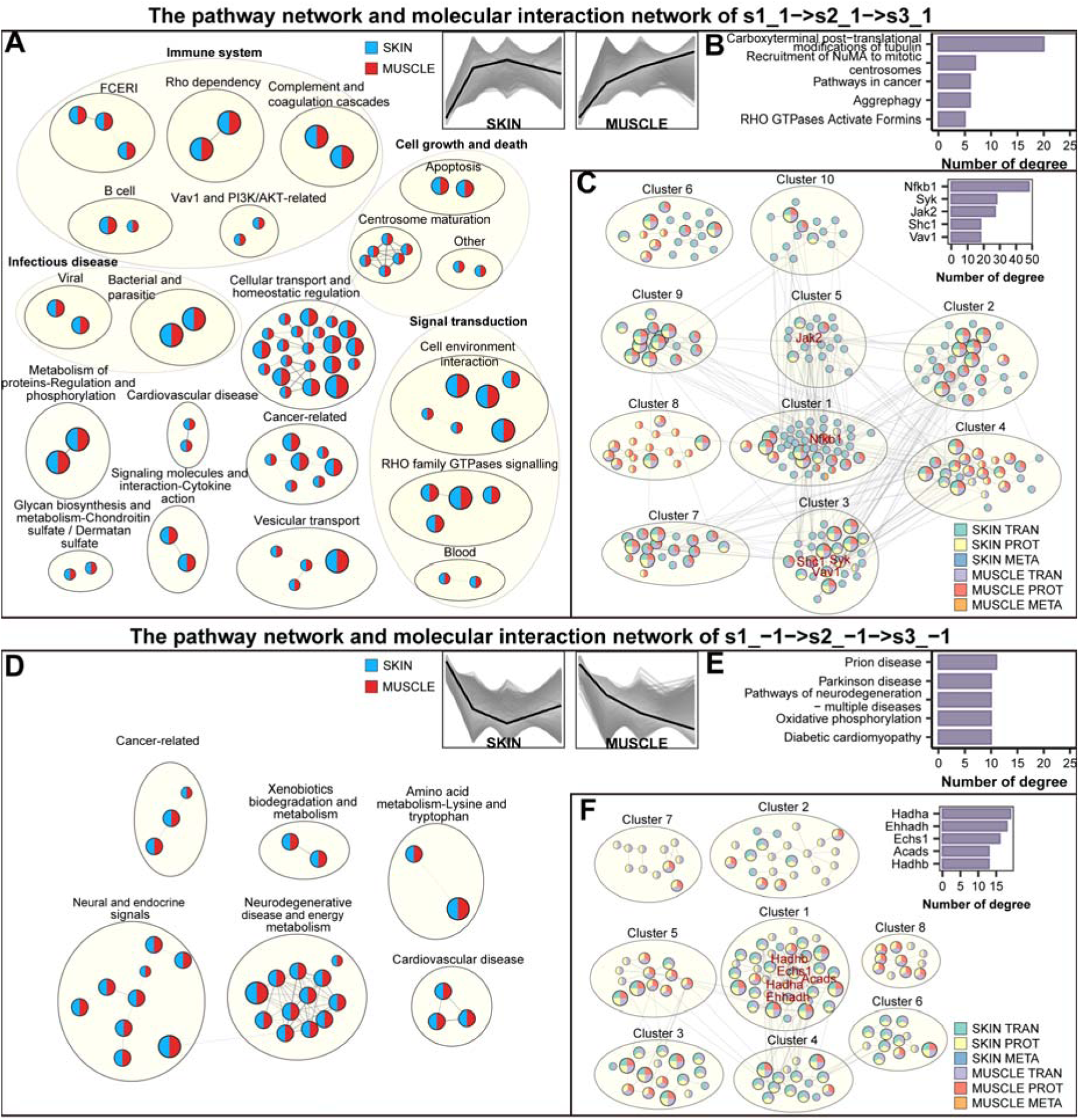
Pathway networks and molecular interactions of the first two major trajectories in skin and muscle. (**A)** Pathway network for pathways jointly enriched in the sustained up-regulation trajectory (s1_1 -> s2_1 -> s3_1) across skin and muscle. Each node represents a pathway; node color indicates source tissue; node size is proportional to the number of tissue and omics categories supporting that pathway. Edges connect pathways with functional similarity (similarity score ≥ 0.5); edge thickness is proportional to similarity. Pathway nodes were clustered using the Louvain algorithm to define biological themes. (**B)** The top five pathway nodes with highest degree (connectivity) in panel **A**. (**C)** Molecular interaction subnetwork of shared upregulated features derived from a large prior-knowledge interaction network. Nodes represent molecules (color indicates tissue and omics category; node size is proportional to the number of tissue and omics categories it encompasses); edges represent high-similarity molecular pairs (similarity score ≥ 0.5); edge thickness is proportional to similarity. Modules were identified by the leading-eigenvector algorithm. (**D–F)** Analogous displays for the sustained down-regulated trajectory (s1_-1 -> s2_-1 -> s3_-1): **D** pathway network; **E** top five high-degree pathway nodes; **F** molecular interaction subnetwork for shared downregulated features.

Skin-specific enrichments emphasized vasculature–immune interactions and carbohydrate metabolism, with “aggregate autophagy” exhibiting high connectivity within the network, suggesting that protein homeostasis regulation plays a crucial role in the skin’s response to compression; skin hubs included Grb2 and Hif1a, pointing to cellular stress responses, hypoxia and angiogenic responses. Muscle-specific enrichments focused on gene-expression regulation, translation, lipid and carbohydrate metabolism, with microtubule-associated modifications also serving as highly connected nodes in the network; muscle hubs included Rac1, Rela, Src and Mapk3, indicating synergistic activation of cytoskeletal reorganization, inflammation, and signal transduction (Supplementary Fig. 11, 13).

#### Sustained down-regulated trajectory (s1_-1 -> s2_-1 -> s3_-1)

Shared downregulated pathways encompassed xenobiotic metabolism, energy metabolism, amino-acid metabolism, mitochondrial and peroxisomal functions, and several pathways linked to neurodegenerative and cardiovascular diseases; prion disease pathways showed high topological centrality (Fig. 5D, E; Supplementary Table 14). Shared high-connectivity molecules included enzymes of fatty-acid β-oxidation and mitochondrial energy metabolism (Hadha, Hadhb, Ehhadh, Echs1, Acads, etc.), clustering in modules enriched for fatty-acid degradation, branched-chain amino-acid catabolism and short-chain fatty-acid metabolism, indicative of coordinated suppression of lipid catabolism and tricarboxylic acid (TCA) cycle-related pathways (Fig. 5F; Supplementary Fig. 13; Supplementary Table 15).

Skin-specific downregulated pathways implicated stress responses, cortisol regulation, lipid and carbohydrate metabolism, with hubs such as Prkacb, Cyp2e1, Adh7, Mapk3 and Rxra, suggesting altered metabolic adaptation and glyco-lipid regulation in skin. Muscle-specific downregulation affected gene expression and repair, insulin signaling, proteostasis and vascular homeostasis; muscle hubs included Prkaca, Mapk1, Mapt, Akt2 and Sod1, reflecting synergistic impairment in cell cycle, mitochondrial, oxidative stress and protein homeostasis (Supplementary Fig. 12, 13).

#### Third trajectory (skin: s1_-1 -> s2_-1 -> s3_0; muscle: s1_0 -> s2_1 -> s3_1)

The third skin trajectory (early decrease then recovery at s3) was enriched for translational control, endoplasmic reticulum (ER) and vesicular transport (e.g., COPI-mediated transport) and mTOR signaling; high-connectivity molecules included Enpp1, Erbb2, Entpd8, Fzd6 and Itga5, highlighting regulation of cell growth, metabolism and adhesion remodeling (Supplementary Fig. 10A–C). The corresponding muscle trajectory (late up-regulation) was enriched for extracellular-matrix metabolism, adhesion and remodeling, immune response and muscle-contraction pathways; muscle hubs included Src, KRAS, Itgav, Ptk2 and Enpp1, emphasizing the importance of adhesion-signal coupling and extracellular matrix (ECM) remodeling in muscle injury progression (Supplementary Fig. 10D–F; Supplementary Fig. 13).

### Immune responses induced by pressure injury

GSEA of immune pathways indicated overall immune activation in both tissues. Skin immune enrichment peaked at s2 and then partially subsided, whereas muscle maintained high immune signals throughout the time course and exhibited a broader repertoire of immune-related changes (Fig. 6A, B; Supplementary Tables 17–18). Integrating the transcriptional and proteomic changes of immune cell marker sets (from [29, 30]; Supplementary Table 21) with pathway enrichment showed skin immune activation concentrated in early–mid stages (s1, s2), while muscle immune activation persisted into mid–late stages (s2, s3) (Supplementary Fig. 14A).

**Fig. 6.**
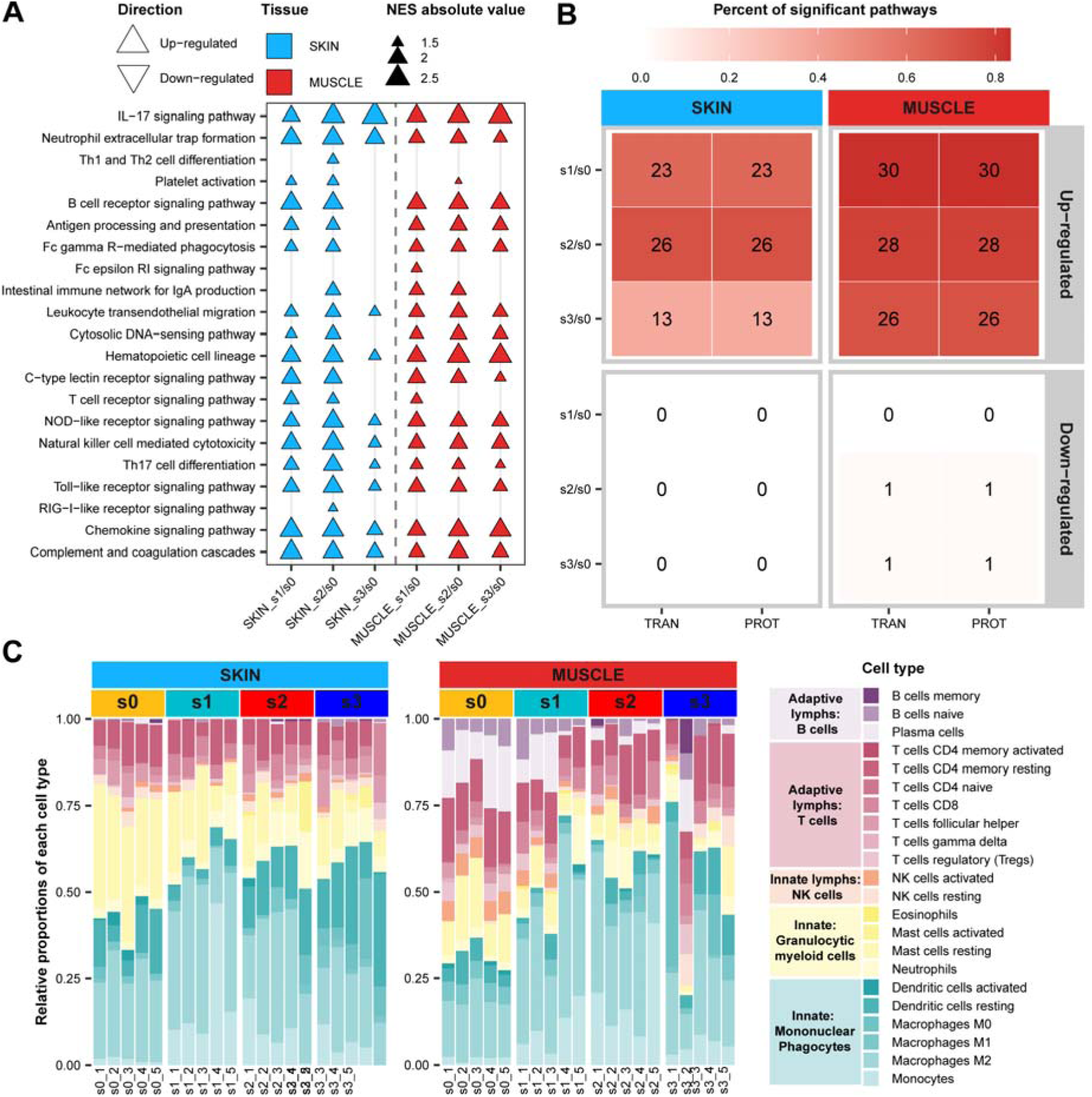
Immune responses induced by pressure injury. (**A)** Gene Set Enrichment Analysis (GSEA) results for KEGG immune-system pathways in skin and muscle across stages; only pathways with FDR < 0.05 are shown. (**B)** Counts of significantly up- or downregulated immune pathways (KEGG and Reactome) per stage for each tissue (FDR < 0.05). (**C)** Immune cell composition inference based on CIBERSORTx: relative abundance of 22 immune cell types in each skin and muscle sample. (each column represents one sample).

CIBERSORTx-based inference of 22 immune cell types across samples revealed that the top three immune cell types by relative abundance in both skin and muscle were monocyte/macrophage, granulocyte/myeloid cells, and T cells. The overall trend showed: as injury progressed, the relative proportion of monocyte/macrophage increased in both tissues; B cells exhibited a relatively high proportion in muscle during the early stage but decreased in the later stage (Fig. 6C). Comparing absolute scores of immune cell types across tissue stages revealed: resting dendritic cells, M2 macrophages, monocytes, neutrophils, and CD8 T cells all significantly increased during injury. Skin reached peak levels primarily in the early-mid stages (s1, s2), while muscle exhibited higher peaks in the mid-late stages (s2, s3). Additionally, skin exhibited distinct differences in immature B cells, M0 macrophages (significantly elevated in stage s3) and quiescent mast cells (gradually decreasing during injury); muscle showed trends toward increased quiescent NK cells and quiescent CD4+ T cells (Supplementary Fig. 14B).

M2 macrophage absolute scores were approximately tenfold higher than M1. Stage-to-stage relative growth rates of M1 and M2 absolute scores showed that in skin both M1 and M2 macrophages increased rapidly during the early injury stage (s1), with M2 increasing faster. However, in the mid–late stages (s2 and s3), M1 growth exceeded M2, suggesting a potential shift toward a pro-inflammatory phenotype in the later course of the disease. Muscle followed a similar pattern: rapid M2 expansion at s1, then relatively greater M1 growth at s2 and minimal change for both thereafter (Supplementary Fig. 14B, C). Using published pro- and anti-inflammatory factor sets [29, 31, 32] (Supplementary Table 22), muscle exhibited a larger number of differential inflammatory factors than skin, with more differentially regulated factors in mid–late stages (s2–s3) than in early stage (s1) for both tissues (Supplementary Fig. 14D, E). Together, multi-omics and deconvolution analyses demonstrate a persistent immune activation and spatiotemporal remodeling of immune cell composition in the injured microenvironment.

### Metabolic alterations induced by pressure injury

Metabolite-class GSEA revealed that skin exhibited upregulation of lipid-related molecules (particularly fatty acyls and glycerides) during injury, while several organic acids and derivatives (carboxylic acids) tended to be downregulated (Fig. 7A, B; Supplementary Table 23). Arachidonic acid (AA), an omega-6 polyunsaturated fatty acid and primary precursor of eicosanoids, whose metabolites function as inflammatory mediators in numerous cellular processes, increased in both skin and muscle (Supplementary Fig. 15A). The detected glycerides in skin were exclusively monoester glycerides, intermediates in lipid metabolism or energy production (Supplementary Fig. 15B). Concomitant downregulation of multiple lipid-related pathways and a significant downregulation of peroxisome pathway proteins at s3 in skin indicate impaired peroxisomal lipid handling (Fig. 7C; Supplementary Fig. 15C, D).

**Fig. 7.**
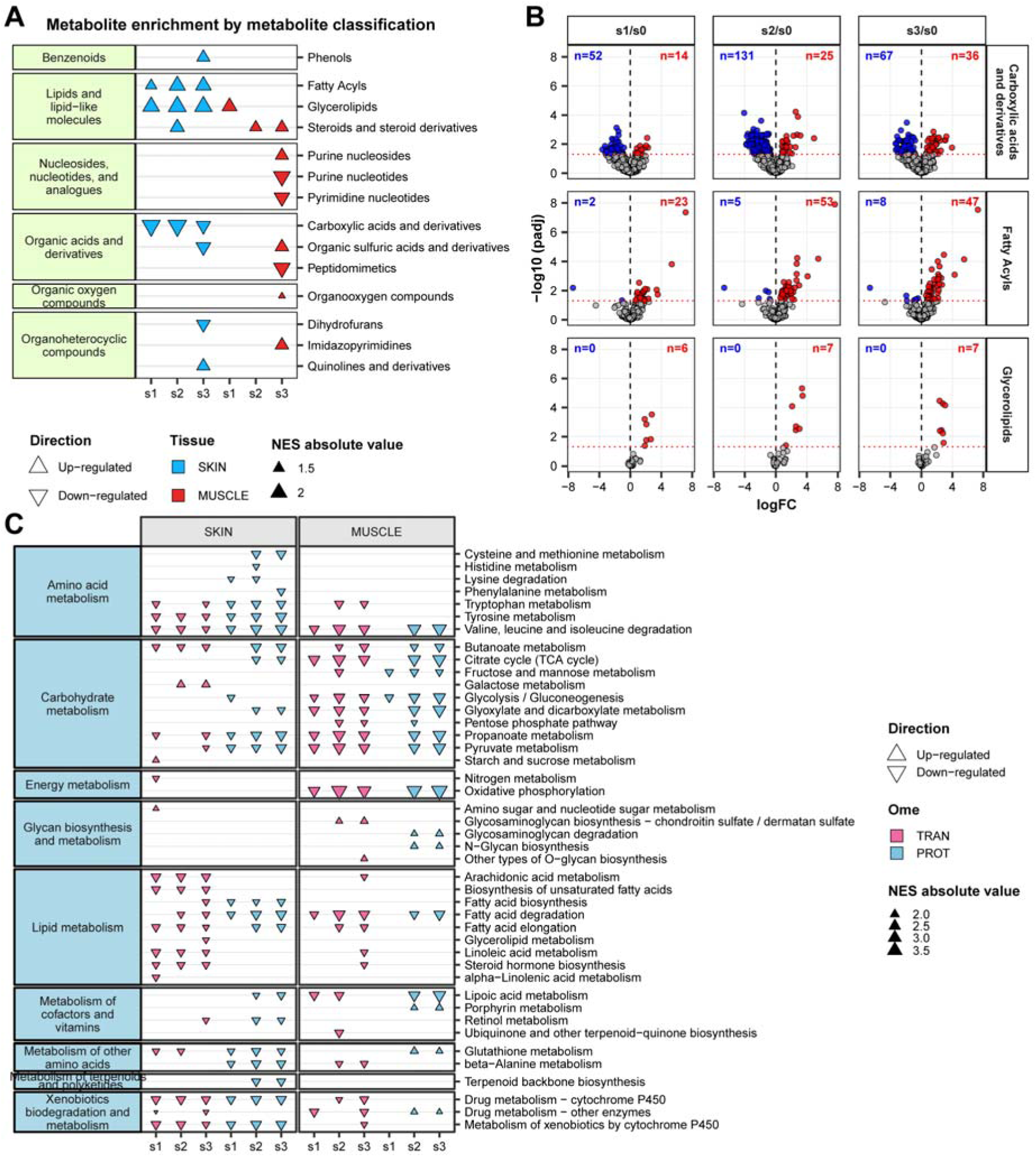
Metabolic response induced by pressure injury. (**A)** Class enrichment analysis (GSEA; FDR < 0.05) of differential metabolites per tissue, based on Human Metabolome Database (HMDB) classification. (**B)** Volcano plots showing metabolite categories with significant changes in skin across stages. Top row: carboxylic acids and derivatives; middle row: fatty acyls; bottom row: glycerolipids. Red points represent upregulated metabolites (FDR < 0.05 and log2FC > 0); blue points represent downregulated metabolites (FDR < 0.05 and log2FC < 0). (**C)** GSEA results (FDR < 0.05) for KEGG metabolic pathway categories in each tissue at each stage.

Skin showed broad downregulation of amino-acid and lipid metabolic pathways across omics, whereas muscle primarily exhibited downregulation of carbohydrate metabolism (Fig. 7C). Proteins associated with the fatty acid degradation pathway were significantly downregulated across all injury stages in skin, consistent with alterations in lipid-related metabolite profiles. TCA cycle pathway was downregulated in both tissues and was particularly pronounced in muscle, suggesting compromised energy metabolism and ATP production (Fig. 7C; Supplementary Fig. 15E).

Based on these findings, we analyzed transcriptional and protein expression changes in 13 mitochondrially encoded genes during muscle injury. Results showed a consistent downregulation trend in identified genes throughout the injury process, indicating dysregulated mitochondrial oxidative phosphorylation pathway activity and resulting energy deficiency (Supplementary Fig. 16A). GSEA using MitoCarta3.0 human mitochondrial gene sets (minimum gene set = 5) revealed pervasive decreases in mitochondrial biogenesis signatures at both transcript and protein levels across muscle injury stages, emphasizing mitochondrial dysfunction during muscle PI progression (Supplementary Fig. 16B; Supplementary Table 24). In summary, integrated metabolomic, transcriptomic and proteomic evidence indicates dysregulation of lipid, carbohydrate and amino-acid metabolism, impairment of TCA and oxidative phosphorylation, and resultant bioenergetic insufficiency that may impede repair and exacerbate disease progression.

### Similarity to human disease

Histological comparison showed concordant features between human PI and the rat model: human PI skin H&E images displayed epidermal hyperplasia, dermal fiber rearrangement and loss of skin appendages; human PI muscle H&E images showed fiber atrophy and fragmentation, and widened inter-fiber spaces. Using the proteomic dataset from Liu et al. [33] after orthologue mapping, log2FC values between mapped human and rat features were significantly positively correlated, with the highest concordance observed at the late stage (s3; Spearman rho = 0.581) (Supplementary Fig. 17A–D). Under a stringent differential threshold (FDR < 0.05 and |log2FC| > 1), Fisher’s exact test showed significant overlap between human and rat differential proteins (two-sided p < 0.05), and an exact binomial test indicated significant directional concordance among overlapping features (two-sided p < 0.05) (Supplementary Fig. 17E, F). Disease-Ontology enrichment of mapped rat features identified pathways linked to human conditions (e.g., immune system diseases, diabetes and acquired metabolic disorders) in both tissues, along with tissue-specific enrichments (e.g., Type-2 diabetes and respiratory-related terms more prominent in skin; cardiomyopathy and muscle disease terms in muscle) (Supplementary Fig. 17G; Supplementary Table 25). These cross-species comparisons support substantial molecular concordance between the rat PI model and human PI and provide a rationale for downstream target validation and translational investigation.

## Discussion

This study employed a longitudinal, multi-tissue (skin and deep muscle) and multi-omics (transcriptome, proteome, metabolome) integrated analysis in a rat pressure injury (PI) model to systematically map the temporal molecular landscape across four key stages of disease progression from baseline to deep muscle exposure. Compared with prior studies limited to a single tissue, omics layer or timepoint, our design captures both temporal resolution and tissue contrasts, enabling stratified characterization of dynamic remodeling in immune, metabolic and signaling networks at gross-pathological, histological and molecular levels. Through multi-omics cross-validation, our data confirm the central involvement of complement and coagulation cascades, neutrophil-mediated responses and mitochondrial oxidative phosphorylation dysfunction, while also revealing pronounced tissue-specific trajectories, early transcriptional programs that initiate downstream responses, and marked inter-individual molecular heterogeneity. Together, these observations generate novel, testable hypotheses for PI pathophysiology (Fig. 8).

**Fig. 8.**
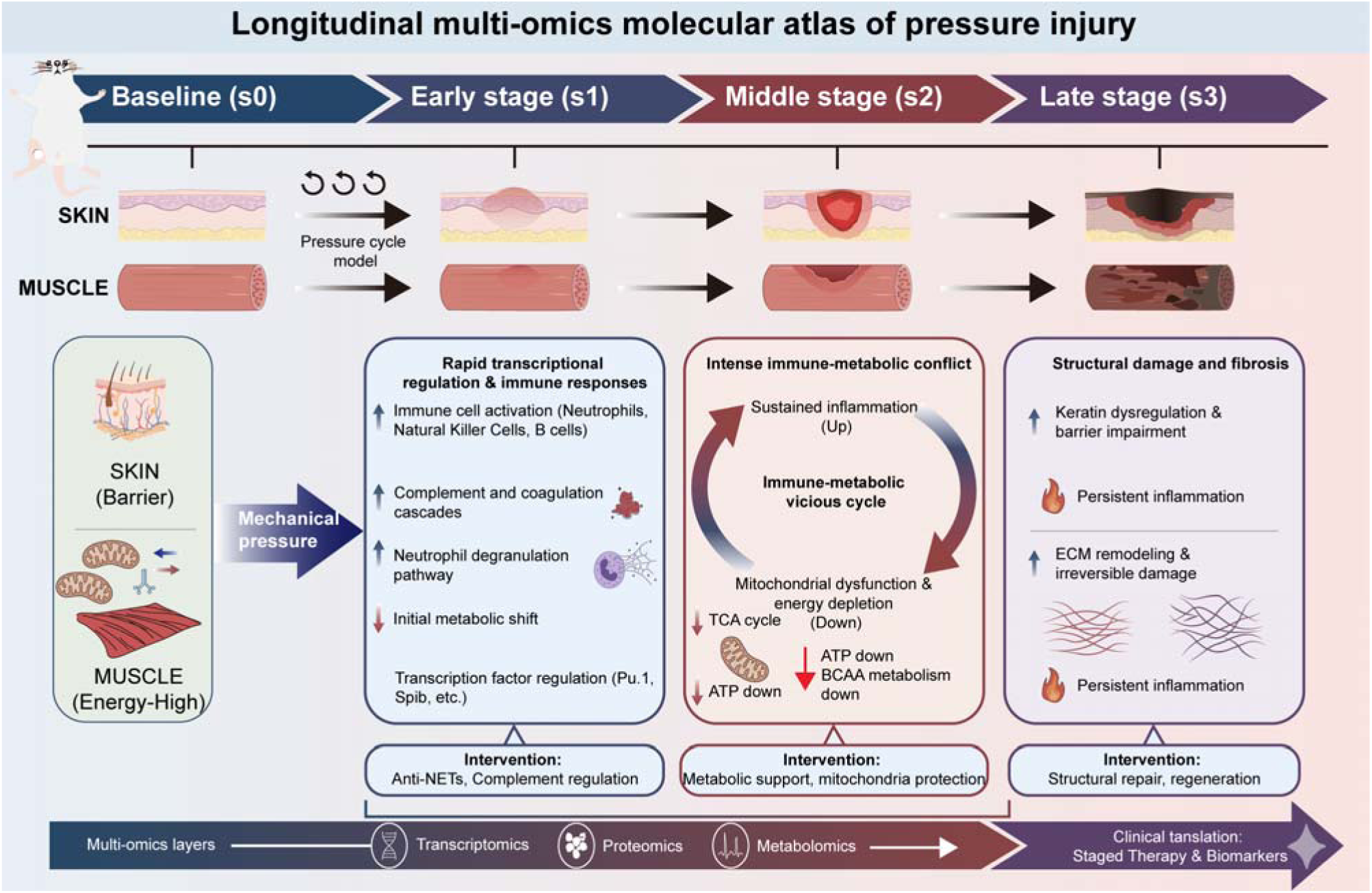
Graphical summary of temporal multi-omics dynamics in pressure injury. This schematic illustrates the stage-resolved molecular landscape of pressure injury across skin and deep muscle based on integrated transcriptomic, proteomic, and metabolomic analyses. Early stages are characterized by rapid activation of transcriptional regulators and innate immune pathways, including complement, coagulation, and neutrophil responses. As injury progresses, a coupled pattern of sustained inflammation and progressive suppression of metabolic and mitochondrial pathways emerges. Skin preferentially engages barrier maintenance, keratinization, and protein homeostasis programs, whereas muscle exhibits pronounced metabolic dysfunction, extracellular matrix remodeling, and structural degradation, contributing to increased susceptibility to irreversible damage. These findings highlight coordinated yet tissue-specific molecular trajectories and suggest stage-dependent therapeutic opportunities targeting immune regulation and metabolic support.

A dominant and persistent signature is activation of the complement system and the coagulation cascade. Complement components (for example C3, C5) and coagulation factors were upregulated from the early stage (s1), with particularly strong signals observed at the protein level. Multiple injury models have demonstrated bidirectional activation and amplification between complement and coagulation (e.g., traumatic brain injury and ischemia–reperfusion [34, 35]), whereby complement cleavage products promote platelet adhesion and coagulation, while thrombin and extracellular matrix alterations can activate complement. This sustained activation may form a vicious positive feedback loop of pro-inflammatory and pro-coagulant effects [36–38]. Concomitantly, we observed rapid early responses in neutrophil-associated pathways, including granule shedding and neutrophil extracellular traps (NETs), accompanied by a concomitant increase in the proportion of infiltrating immune cells. Although NETs can trap pathogens and clear debris, excessive or sustained NETosis damages epithelial and endothelial barriers, promotes immunothrombosis, impairing microcirculation and delaying repair [39]. These findings suggest that among complement, coagulation, and NETs crosstalk may form a central, self-amplifying axis in the PI microenvironment that sustains chronic inflammation and microcirculatory dysfunction; stage-targeted modulation of NET formation or local complement activity may therefore represent a potential therapeutic strategy to mitigate persistent inflammation and improve healing.

Disordered energy metabolism emerged as a core feature in the progression of muscle injury. We observed coordinated downregulation of fatty-acid β-oxidation, the tricarboxylic-acid (TCA) cycle and oxidative phosphorylation at transcript and protein levels. Mitochondrial dysfunction is a well-established driver of injury in ischemia–reperfusion and other organ injuries, reducing ATP production, increasing reactive oxygen species (ROS) and activating cell-death pathways that impede repair [40–42]. In network reconstructions, sustained upregulated immune hubs (Nfkb1, Jak2, Syk, Vav1) contrasted with a sustained downregulated module enriched for mitochondrial and energy metabolic factors (for example Hadha). This coupled pattern of “immune up-regulation — metabolic down-regulation” suggests a biological trade-off involving resource reallocation: the body prioritizes energy mobilization for inflammatory clearance and barrier maintenance during early injury, but under persistent mechanical stress mitochondrial impairment undermines energy supply and reparative capacity, promoting irreversible tissue damage [43, 44]. Suppression of branched-chain amino-acid (BCAA) metabolism in both tissues further supports impaired anaplerosis to the TCA and suggests nutrient-metabolic interventions (for example BCAA supplementation or metabolic pathway modulation) as candidate therapeutic strategies [44, 45]. The more severe collapse of TCA and oxidative phosphorylation in muscle provides a molecular rationale for the clinical observation that deep muscle is more susceptible to irreversible injury under comparable external load [46, 47].

The immediate response to mechanical stress is rapidly initiated through gene regulatory networks dominated by transcription factors such as NF-κB, ETS family members (e.g., PU.1), KLF, and HIF. These TFs regulate genes encoding chemokines, adhesion molecules and complement components and drive recruitment and activation of innate immune cells [48, 49]. For example, PU.1 directly regulates specific chemokine genes (e.g., Ccl22) and participates in the functional programming of macrophages and dendritic cells [50]. Notably, microtubule post-translational modifications (PTMs) (for example detyrosination, acetylation) emerge as highly connected nodes in sustained upregulated trajectories. As the “highways” of the cytoskeleton, the PTM status of microtubules directly regulates intracellular transport (including inflammasomes and lysosomes) and the migration and polarization of immune cells [51, 52]. This discovery links cellular mechanical responses to immune inflammation: mechanical stress may promote intracellular transport and secretion of inflammatory mediators by altering microtubule PTMs, thereby exacerbating tissue damage. This suggests that stabilizing microtubule dynamics may help attenuate mechanically induced inflammatory amplification.

Tissue-specific responses were pronounced. Skin mounted an adaptive program centered on barrier maintenance and proteostasis: keratin family members (Krt18, Krt8) and cornified-envelope components were strongly induced, and the aggrephagy pathway and associated molecules (e.g., Grb2, Hif1a) were engaged, indicating the skin initiated a potent protein quality control mechanism, attempting to clear misfolded proteins produced by oxidative stress and maintain cellular survival [53, 54]. Such defense likely underlies why early stage (s1) lesions may remain non-ulcerative. By contrast, muscle exhibited greater structural disassembly and metabolic collapse: mitochondrial dysfunction, impaired oxidative phosphorylation and remodeling of ECM and adhesion pathways render muscle more vulnerable to irreversible necrosis and fibrosis. These tissue-dependent difference reflect differing structural and energetic demands: skin, as the primary barrier, prioritizes restoring barrier integrity, whereas muscle, a high-energy structural tissue, relies more heavily on energy supply and cytoskeletal integrity. Once mitochondrial function is compromised, muscle is more susceptible to irreversible structural damage, rapidly progressing toward necrosis and fibrosis. This finding underscores that reliance on cutaneous appearance alone may miss deep tissue compromise. Clinically, this supports the need for deeper metabolic monitoring and tissue-specific, stage-adapted interventions [6, 53].

The temporal molecular map suggests stage-specific therapeutic strategies. The early stage (s1) represents a “pre-reaction period” characterized by transcriptional regulation and immune recruitment. At this stage, tissue has not yet undergone irreversible necrosis. Targeting overactivated immune pathways (e.g., inhibiting NET formation, locally modulating complement activity) may help attenuate inflammatory amplification. Mid-stage (s2) is characterized by intense immune–metabolic cross-talk; combined anti-inflammatory and metabolic support (for example PPAR agonists, mitochondria-protective agents or targeted nutritional approaches) may help restore metabolic homeostasis and limit fibrotic progression. The late stage (s3) is dominated by structural loss and fibrosis, where surgical intervention and promotion of tissue regeneration are likely required. Existing literature indicates that PPARγ agonists can modulate macrophage phenotypes and promote repair, supporting the potential of pathway interventions in repair [55, 56]. Our high-variability feature set further provides candidate biomarkers for stratified medicine — prioritizing early, protein-detectable molecules (e.g., complement components, inflammatory mediators or ECM degradation factors) may accelerate translational validation.

Despite the strengths of this study in multi-omics integration and temporal resolution, limitations remain. First, despite histological and molecular concordance between the rat model and human PI, anatomical and immunological differences (e.g., presence of the panniculus carnosus) limit direct clinical extrapolation; therefore, key pathways and biomarkers require validation in models closer to humans or in clinical samples [57, 58]. Second, bulk omics approaches average signals across heterogeneous cell populations and may obscure cell-type specific dynamics; integration with single-cell and spatial omics is needed to resolve the cellular drivers and spatiotemporal interactions. Third, candidate mechanisms identified in this study (e.g., NETs, microtubule PTMs) require direct functional testing by genetic or pharmacological perturbation to establish causality.

## Conclusion

In conclusion, this longitudinal multi-omics atlas of PI across skin and deep muscle reveals an early transcriptional program that drives immune recruitment and activation of complement and coagulation pathways, coupled with progressive metabolic and mitochondrial dysfunction in muscle and protein homeostasis responses in skin. The resulting atlas generates experimentally tractable hypotheses and provides a molecular rationale for stage- and tissue-specific interventions and biomarker development for PI.

## Abbreviations

PI: Pressure injury
HAPIs: Hospital-acquired pressure injuries
NPUAP: National Pressure Ulcer Advisory Panel
H&E: Hematoxylin–eosin
VST: Variance-stabilizing transformation
CPM: Counts per million
CV: Coefficient of variation
KNN: K-nearest neighbors
QC: Quality control
PCA: Principal component analysis
t-SNE: T-distributed stochastic neighbor embedding
RIN: RNA integrity number
FDR: False discovery rate
GSEA: Gene set enrichment analysis
EM: Expectation-Maximization
TPM: Transcripts per million
TAF: Tissue-stained area fraction
CVF: Collagen volume fraction
TF: Transcription factor
Q-Q: Quantile-quantile
TCA: Tricarboxylic acid
ECM: Extracellular matrix
AA: Arachidonic acid
NETs: Neutrophil extracellular traps
ROS: Reactive oxygen species
BCAA: Branched-chain amino-acid
PTMs: Post-translational modifications

## Supplementary information

Supplementary Material 1: Supplemental Methods, Supplemental Results and Supplemental Figures S1-S17. Supplementary Material 2: Supplemental Tables S1-S25.

## Acknowledgements

We thank for the support by the National Natural Science Foundation of China (No. 12272251).

## Author contributions

Conceptualization: FJ, MWA, XJ. Methodology: FJ, LYL, YTZ, TZ, XL, JCW, NXL, YRJ, MWA, XJ. Software: FJ, XL. Investigation: LYL, YTZ, TZ, XNL. Resources: MWA, XJ. Funding acquisition: MWA. Supervision: MWA. Writing—original draft: FJ. Writing—review & editing: MWA, XJ. All the authors reviewed and approved the final version of the manuscript.

## Data availability

The raw sequencing data generated in this study have been deposited in appropriate public repositories: RNA-seq data have been deposited in the Sequence Read Archive (SRA) of the NCBI under BioProject accession PRJNA1391872; mass spectrometry proteomics data have been deposited in the ProteomeXchange Consortium via the iProX partner repository under accession PXD072436; untargeted metabolomics data have been deposited in MetaboLights under accession MTBLS13566. External reference resources used in analyses are publicly accessible as follows: Ensembl rattus norvegicus, mRatBN7.2 genome (https://ftp.ensembl.org/pub/release-110/fasta/rattus_norvegicus/dna/Rattus_norvegicus.mRatBN7.2.dna.toplevel.fa.gz) and gene annotation files (https://ftp.ensembl.org/pub/release-110/gtf/rattus_norvegicus/Rattus_norvegicus.mRatBN7.2.110.gtf.gz); UniProt rat protein reference (https://www.uniprot.org/uniprotkb?query=10116&facets=model_organism%3A10116, accessed 26/6/2024); RGD rat gene annotation file (https://download.rgd.mcw.edu/data_release/RAT/GENES_RAT.txt, accessed 1/6/2025); Reactome pathway collection (https://reactome.org/download/current, accessed 1/8/2025); Reactome molecular interactions (https://download.reactome.org/93/interactors/reactome.all_species.interactions.psi-mitab.txt, accessed 24/8/2025); Rat–human orthologue mappings (https://download.rgd.mcw.edu/data_release/RAT/ORTHOLOGS_RAT.txt, accessed 16/9/2025); Mitochondria-associated genes and pathways MitoCarta3.0 (https://www.broadinstitute.org/mitocarta/mitocarta30-inventory-mammalian-mitochondrial-proteins-and-pathways); CIBERSORT LM22 immune cell gene signature matrix (https://doi.org/10.1038/nmeth.3337).

## Code availability

No novel code or algorithms were developed for this study. All analyses can be reproduced using the software, packages and workflows described in the Methods. Reasonable requests for scripts, pipelines or additional implementation details may be directed to the corresponding author.

## Declarations

### Ethics approval and consent to participate

Not applicable.

### Consent for publication

Not applicable.

