## Supplementary Material1 for "A Cross-tissue Temporal Multi-omics Atlas Reveals the Molecular Architecture of Pressure Injury"

### Supplemental Methods

#### Histopathological quantification

Quantitative metrics were defined as follows: (1) Epidermal thickness: for each H&E image (10×), five random locations were measured and the mean reported; (2) Collagen volume fraction (CVF): for each Masson image (40×), five random fields were selected, color deconvolution applied, and CVF calculated as CVF = Blue area / (Blue area + Red area); (3) Mean muscle fiber cross-sectional area (CSA): for each Masson image (40×), five random fields were selected and CSA calculated as CSA = Total cross-sectional area of fibers / Number of fibers, mean reported; (4) Tissue-stained area fraction (TAF): for each Masson image (40×), five random fields were selected, color deconvolution applied, and TAF calculated as TAF = (Blue area + Red area) / Total image area. Image analysis was performed in Fiji (ImageJ). Multi-group comparisons used one-way ANOVA with Tukey HSD post-hoc test. Significance was set at two-sided p < 0.05.

#### Transcriptome sequencing

Total RNA was extracted using TRNzol Universal (TIANGEN) and RNA integrity assessed via the Agilent 5400. Libraries were prepared with the Fast RNA-seq Lib Prep Kit V2 (ABclonal, Cat. RK20306). mRNA was enriched using Oligo (dT) magnetic beads, fragmented, and synthesized into first-strand cDNA using random hexamer primers, followed by second-strand cDNA synthesis. Libraries were completed by end repair, A-tailing, adapter ligation, fragment selection, amplification and purification. Library quantification and quality control were performed using Qubit 2.0 Fluorometer, real-time quantitative PCR (q-PCR) and a bioanalyzer, respectively. Paired-end sequencing (PE150) was carried out on the Illumina NovaSeq platform.

#### Proteome sequencing

Tissue samples were ground and proteins extracted in DB lysis buffer (6 M urea, 100 mM TEAB, pH 8.5) to a final volume of 100 µL. Samples were digested with trypsin in 100 mM TEAB at 37°C for 4 h. Formic acid was added to adjust pH < 3 and samples were centrifuged at 12,000 g for 5 min at room temperature. Supernatants were desalted on C18 cartridges, washed (0.1% formic acid, 3% acetonitrile) three times and eluted (0.1% formic acid, 70% acetonitrile); eluates were lyophilized for storage. Peptides were separated on a Vanquish™ Neo UHPLC (Thermo Fisher) with a C18 pre-column (174500, 5 mm × 300 µm, 5 µm) and a C18 analytical column (ES906, PepMap™ Neo, 150 µm × 15 cm, 2 µm) at 50°C. Mass spectrometry was performed on an Orbitrap Astral (Thermo Fisher) with an Easy-spray ESI source (spray voltage 2.0 kV, ion transfer tube temperature 290°C). Data were acquired in data-independent acquisition (DIA) mode: full MS1 scan range m/z 380–980 at resolution 240,000, AGC target 500%, max injection time 3 ms; 300 DIA windows, 2 Th window width, NCE 25%; MS2 range m/z 150–2000 at Astral resolution 80,000.

#### Non-targeted metabolomics sequencing

Tissue samples were ground in liquid nitrogen for metabolite extraction and analyzed by LC-MS on a Q Exactive™ HF-X mass spectrometer (Thermo Fisher) coupled to a Vanquish UHPLC. Chromatography separation used a Hypersil Gold column. MS scan range was m/z 100–1500. ESI settings: spray voltage 3.5 kV, sheath gas flow rate 35 psi, aux Gas flow rate 10 L/min, capillary temperature 320°C, S-lens RF level 60, aus gas heater temperature 350°C. Data-dependent scanning (DDA) was performed in both positive and negative ion modes to acquire MS/MS spectra. Quality control (QC) samples were prepared by pooling equal volumes of all experimental samples to monitor system stability; blanks samples (53% methanol) were used to remove background noise.

### Supplemental Results

#### Quality control of omics data

Transcriptome, proteome, and non-targeted metabolome sequencing were performed on skin and muscle samples (sequencing was conducted on subsets of all samples; see Supplementary Fig. 1B and the Methods for the detailed workflow). The median RNA integrity number (RIN) for transcriptome samples across all groups exceeded 6, indicating that the RNA quality was suitable for downstream analysis (Supplementary Fig. 2A, E; Supplementary Table 2). No obvious outlier samples were detected across any omics modality (Supplementary Fig. 2B, F). Furthermore, despite the presence of samples processed across different batches in the transcriptome analysis, batch-effect assessment revealed no apparent systematic bias (Supplementary Fig. 2C, G). Principal component analysis (PCA) and t-distributed stochastic neighbor embedding (t-SNE) demonstrated good separation between groups and tight within-group clustering across omics (Fig. 2A, G; Supplementary Fig. 2D, H). Metabolomics quality control (QC) samples had correlation coefficients > 0.99 in both positive and negative ion modes, indicating robust instrument stability (Supplementary Fig. 2I, K).

After filtering we detected 16,306 transcripts (skin 16,039; muscle 15,190), 9,054 proteins (skin 8,708; muscle 6,597) and 2,923 metabolites (skin 2,923; muscle 2,923) across the study (Fig. 1A). Analysis of molecules expressed as non-zero across all samples showed that the number of transcripts in muscle increased progressively with injury, and that protein counts in experimental groups exceeded baseline (Supplementary Fig. 2J, L). Overall, all omics datasets passed necessary quality controls and were deemed suitable for downstream analyses.

### Supplemental Figures

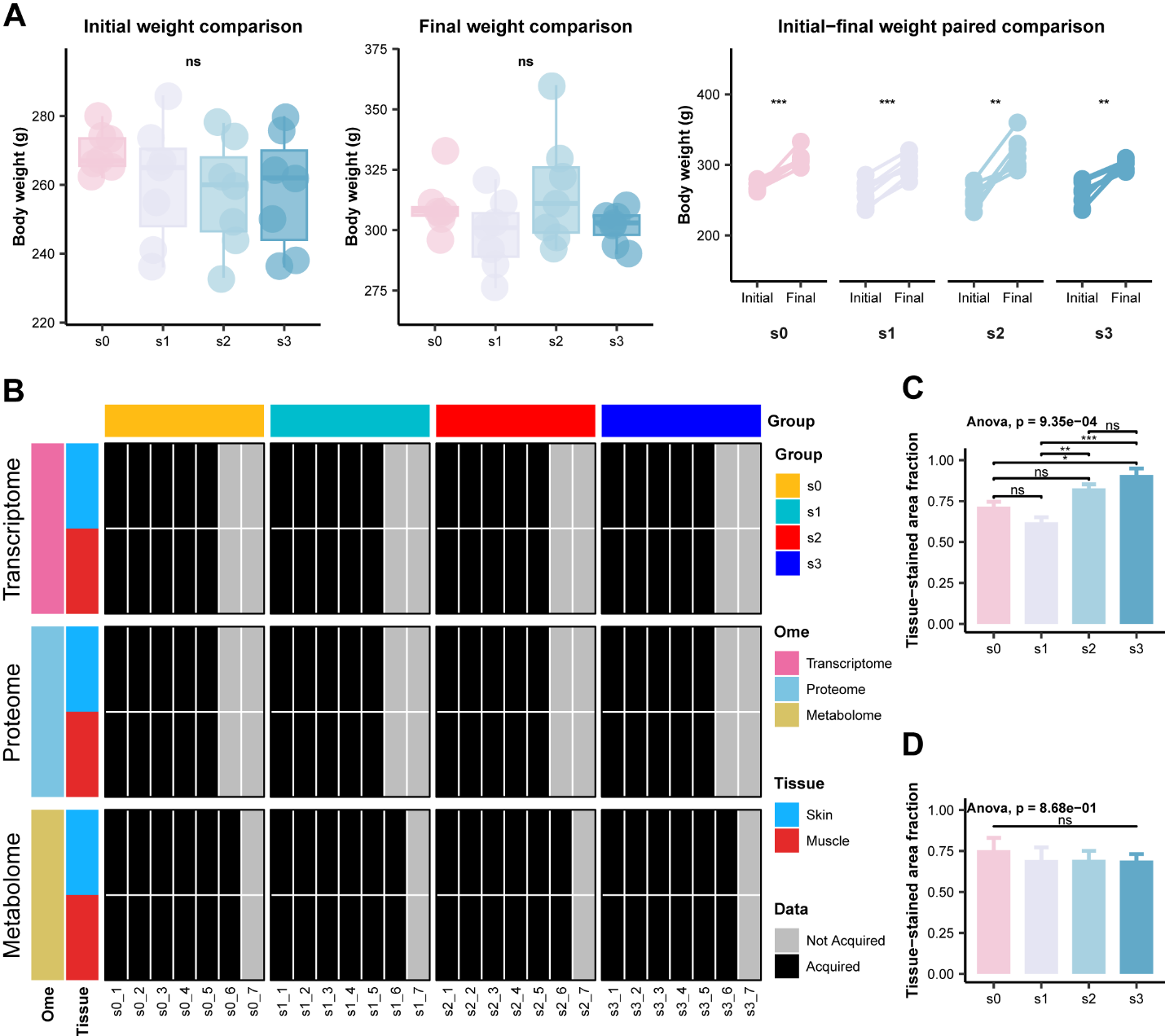

**Fig. S1. Additional animal-phenotype and study-design information.** (**A**) Body-weight comparisons (n = 7 per group). Left, initial body weight after randomization (eight-week-old rats); center, final body weight by group at study endpoint; right, within-group paired comparison of initial versus final body weight. (**B**) Sample availability across omics platforms: number of available samples per group and per platform. (**C**) Skin tissue area fraction (TAF) by group (n = 3). (**D**) Muscle tissue area fraction (TAF) by group (n = 3). Data in c and d are presented as mean ± SEM. Significance markers: ns, p ≥ 0.05; *p < 0.05; **p < 0.01; ***p < 0.001.

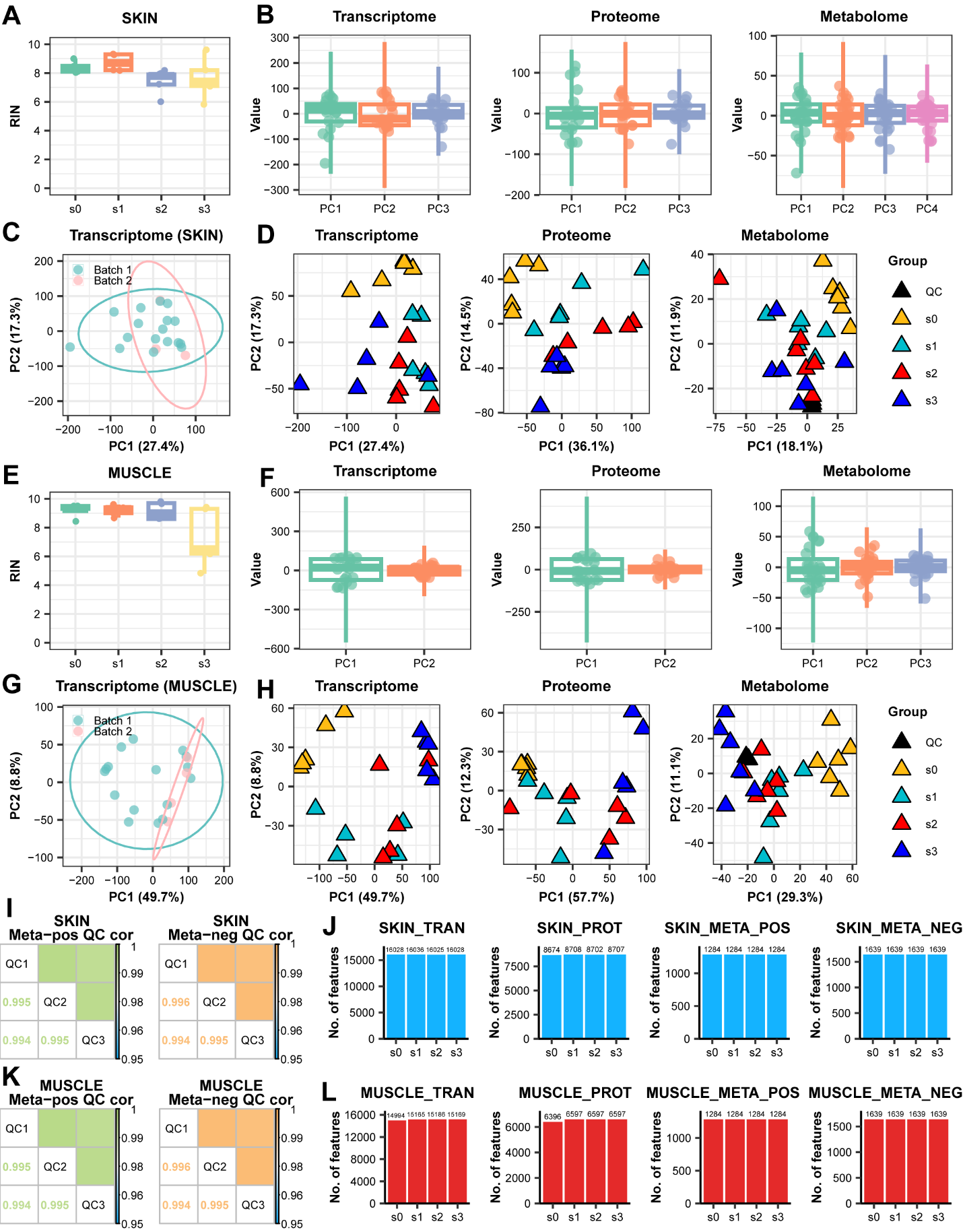

**Fig. S2. Omics data quality control.** (**A**) Boxplots of RNA integrity number (RIN) distributions for skin samples (per-group RINs). (**B**) Outlier detection for skin samples. (**C**) PCA of skin transcriptomes colored by batch (batch-effect assessment). (**D**) PCA of all skin omics samples (each point represents one sample); metabolomics QC is the pooled QC sample from all experimental samples. (**E**) Boxplots of RIN distributions for muscle samples. (**F**) Outlier detection for muscle samples. (**G**) PCA of muscle transcriptomes (batch assessment). (**H**) PCA of all muscle omics samples. (**I**) Pearson correlations among skin metabolomics QC samples in positive and negative ion modes. (**J**) Numbers of features detected (“expressed in at least one sample”) per omics and stage for skin. (**K**) Pearson correlations among muscle metabolomics QC samples in positive and negative ion modes. (**L**) Numbers of features detected per omics and stage for muscle. Boxplot annotations in a and e: lower edge represents the first quartile (Q1), center line represents the median, upper edge represents the third quartile (Q3); whiskers denote non-outlier minimum and maximum. Box plot whiskers in b and f are calculated as Q3 + 3×IQR and Q1 − 3×IQR (IQR represents interquartile range).

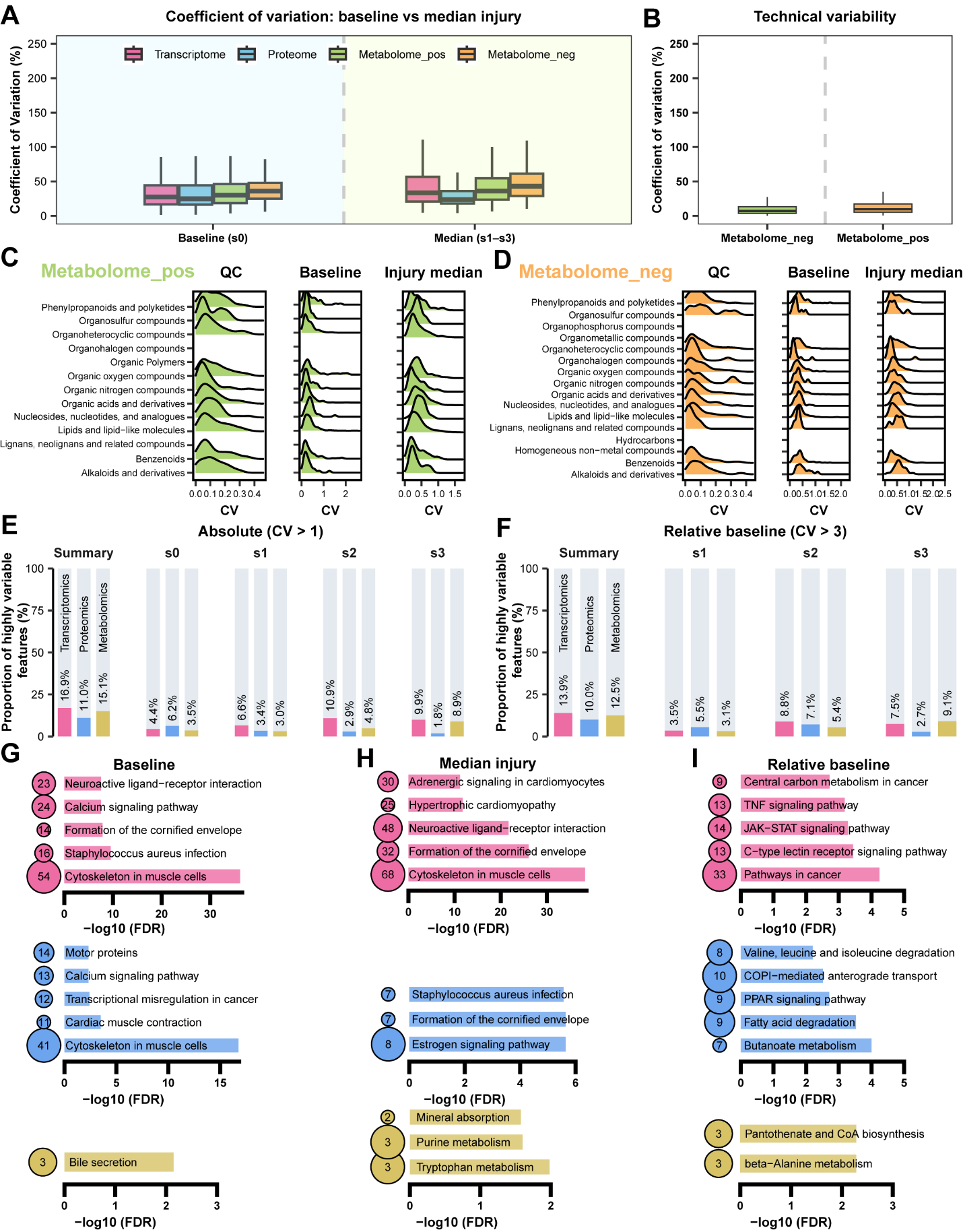

**Fig. S3. Molecular variability in skin.** (**A**) Boxplots comparing coefficient of variation (CV) across omics groups at baseline and post-injury. (**B**) Distribution of CVs for metabolomics QC samples (positive and negative ion modes). (**C**) CV distribution of HMDB-classified metabolites (positive ion mode) in QC, baseline and post-injury samples. (**D**) CV distribution of HMDB-classified metabolites (negative ion mode) in QC, baseline and post-injury samples. (**E**) Counts of high-variability features (CV > 100%) per omics and stage. (**F**) Counts of features with injury-to-baseline CV ratio > 300% per omics and stage. (**G**–**I**) Pathway enrichment (top five) for high-variability features under three conditions: baseline, post-injury median and relative-to-baseline (FDR < 0.05). Top: transcriptome; middle: proteome; bottom: metabolome. Boxplot annotations: lower edge represents the first quartile (Q1), center line represents the median, upper edge represents the third quartile (Q3); whiskers denote non-outlier minimum and maximum.

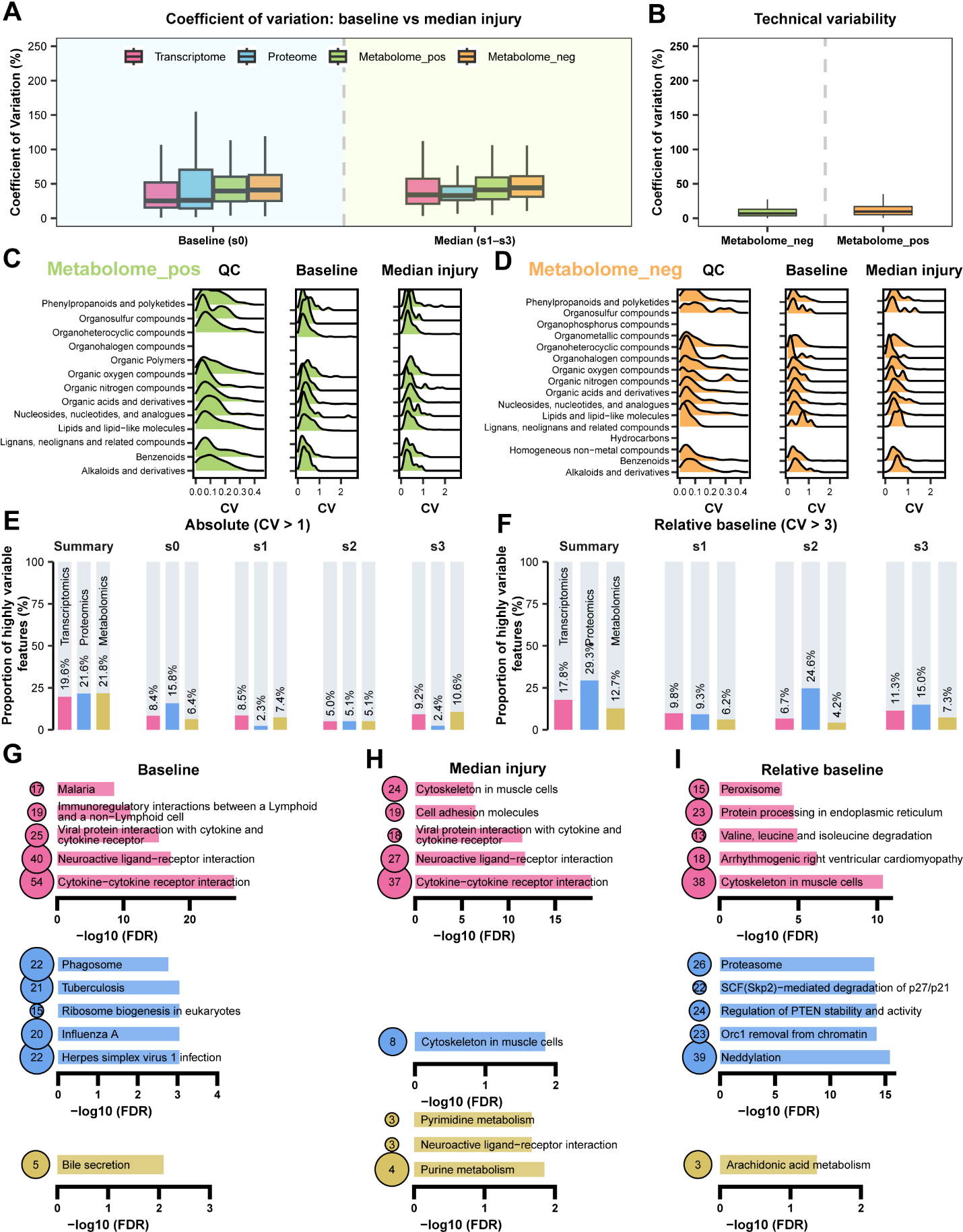

**Fig. S4. Molecular variability in muscle.** (**A**) Boxplots comparing coefficient of variation (CV) across omics groups at baseline and post-injury. (**B**) Distribution of CVs for metabolomics QC samples (positive and negative ion modes). (**C**) CV distribution of HMDB-classified metabolites (positive ion mode) in QC, baseline and post-injury samples. (**D**) CV distribution of HMDB-classified metabolites (negative ion mode) in QC, baseline and post-injury samples. (**E**) Counts of high-variability features (CV > 100%) per omics and stage. (**F**) Counts of features with injury-to-baseline CV ratio > 300% per omics and stage. (**G**–**I**) Pathway enrichment (top five) for high-variability features under three conditions: baseline, post-injury median and relative-to-baseline (FDR < 0.05). Top: transcriptome; middle: proteome; bottom: metabolome. Boxplot annotations as in **Fig. S3**.

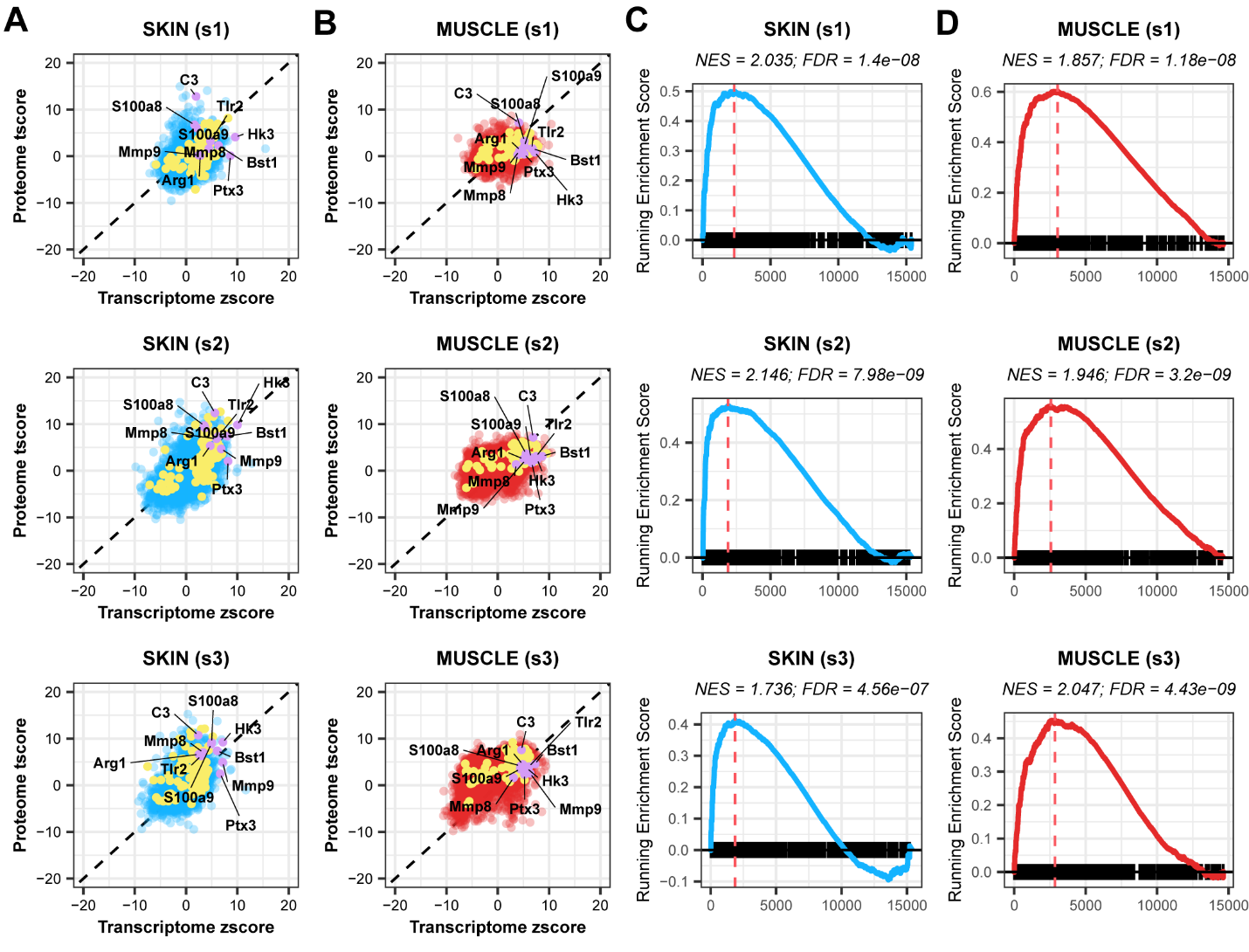

**Fig. S5. Molecular expression and GSEA results for the neutrophil degranulation pathway.** (**A**) Skin: overview of transcripts and proteins within the neutrophil degranulation pathway that are jointly differential across stages (transcript z-score and protein t-score summaries). (**B**) Muscle: corresponding transcript and protein joint-differential summaries. (**C**) GSEA results for the neutrophil degranulation pathway across skin stages. (**D**) GSEA results for the neutrophil degranulation pathway across muscle stages. Color legend in **A** and **B**: blue/red dots represent shared transcription-protein molecules; yellow dots represent genes differentially expressed at both transcription and protein levels within the pathway; purple dots highlight the top ten genes ranked by log2FC at the transcription level across all tissues (i.e., hallmark genes exhibiting substantial changes under multiple conditions).

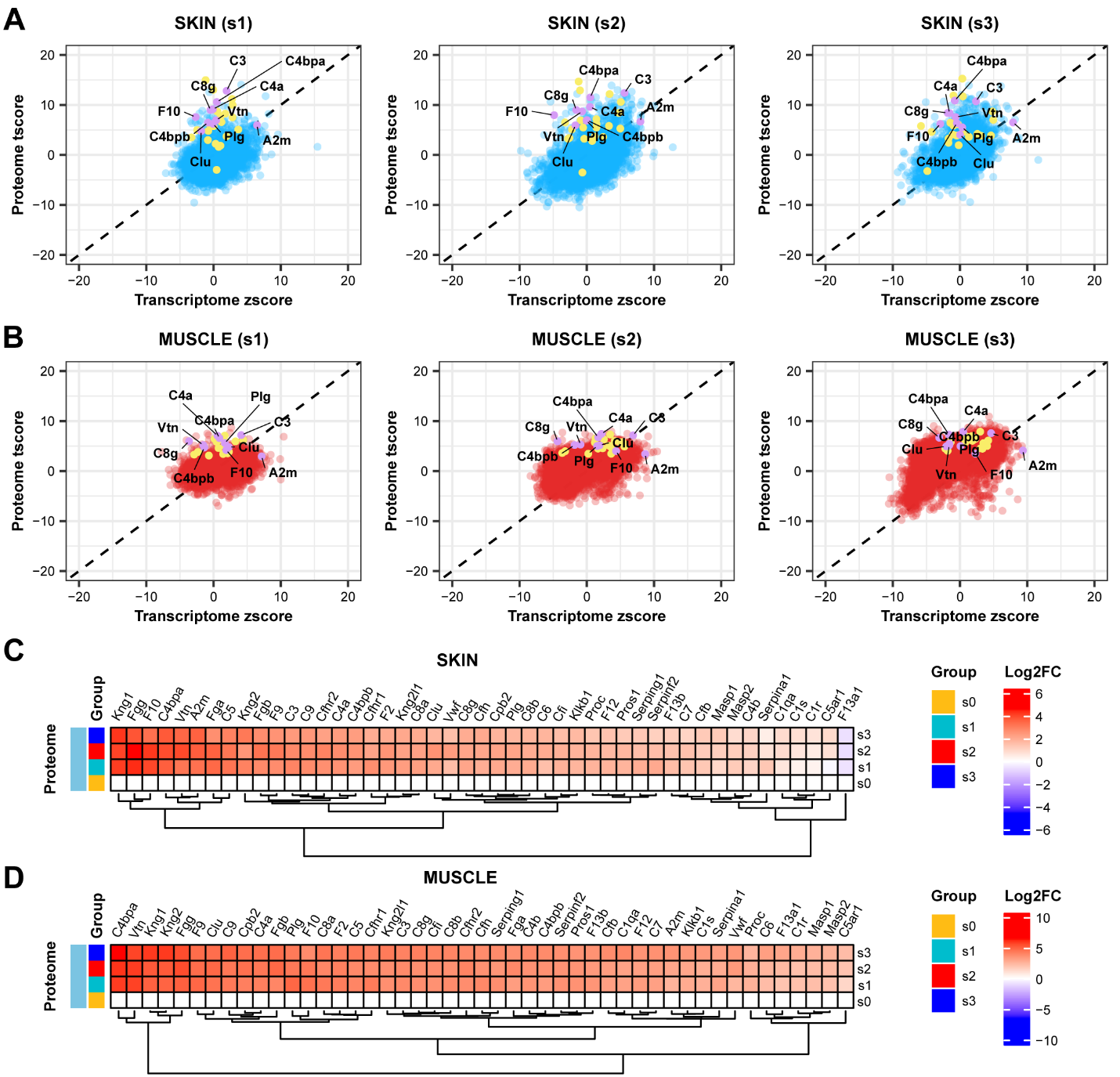

**Fig. S6. Overview of molecular expression in complement and coagulation cascade pathway.** (**A**) Skin: overview of transcripts and proteins within the complement and coagulation cascade pathway that are jointly differential across stages (transcript z-score and protein t-score summaries). (**B**) Muscle: corresponding joint summary. (**C**) Skin: distribution of protein log2FC for complement and coagulation cascade pathway members across stages. (**D**) Muscle: corresponding protein log2FC distribution. Color legend as in **Fig. S5**.

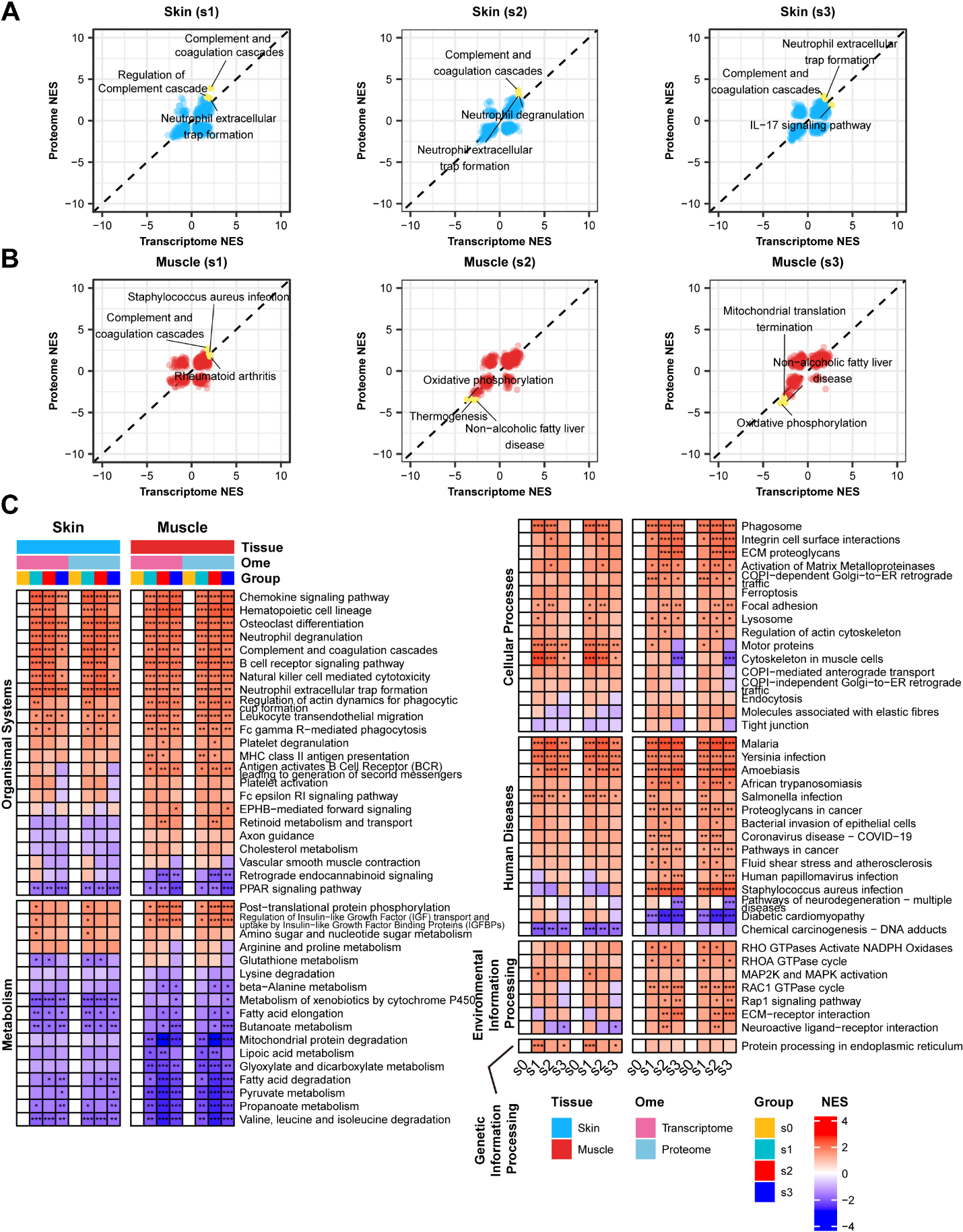

**Fig. S7. GSEA results for transcriptome and proteome pathways across tissues.** (**A**) Skin: scatter plot of normalized enrichment scores (NES) for transcripts and proteins across stages (each point represents a pathway/stage combination); the three highest NES shared transcript–protein pathways are highlighted (yellow). (**B**) Muscle: corresponding NES scatter plot with the three highest shared pathways highlighted. (**C**) Heatmap of NES for pathways jointly enriched in transcriptome and proteome across tissues and stages (shared pathways). Color legend for **A** and **B**: blue/red indicate pathways detected in both transcript and protein layers. Significance annotations: *p < 0.05; **p < 0.01; ***p < 0.001.

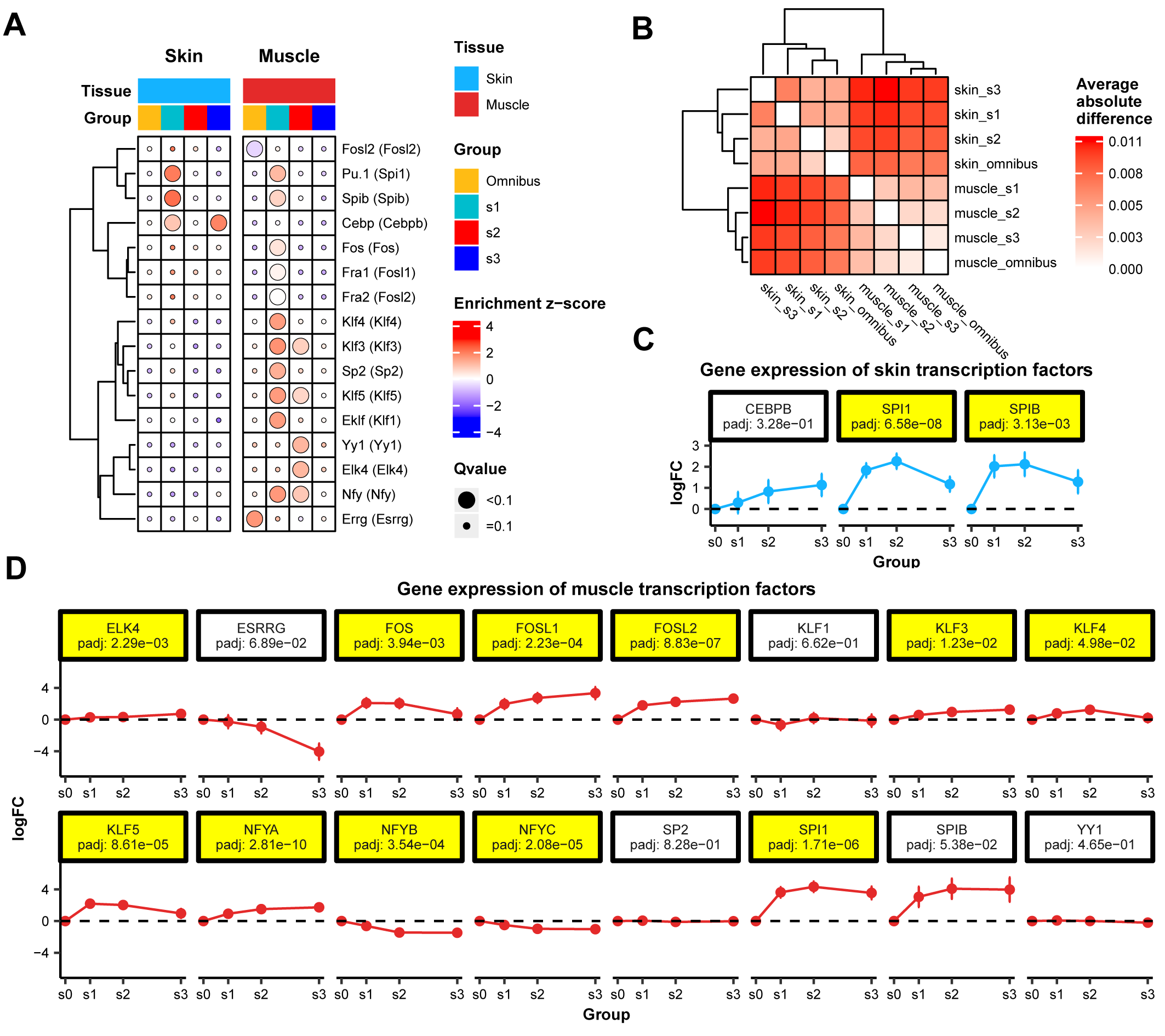

**Fig. S8. Transcription factor (TF) motif enrichment among differential transcripts.** (**A**) Heatmap of TF motif enrichment (z-score) across overall and stage-specific differential transcript sets. Only TFs with q < 0.1 are shown; TFs are hierarchically clustered by their enrichment scores across groups. (**B**) Heatmap of average absolute differences in motif enrichment scores between tissues (hierarchical clustering), illustrating inter-tissue divergence in TF enrichment patterns. (**C**) Line plots of log2FC (with lfcSE error bars) across groups for TFs significantly enriched in panel **A** (skin). s0 is set to log2FC = 0. Padj denotes the adjusted p value for the overall comparison. (**D**) Corresponding TF log2FC line plots for muscle (annotation as in **C**).

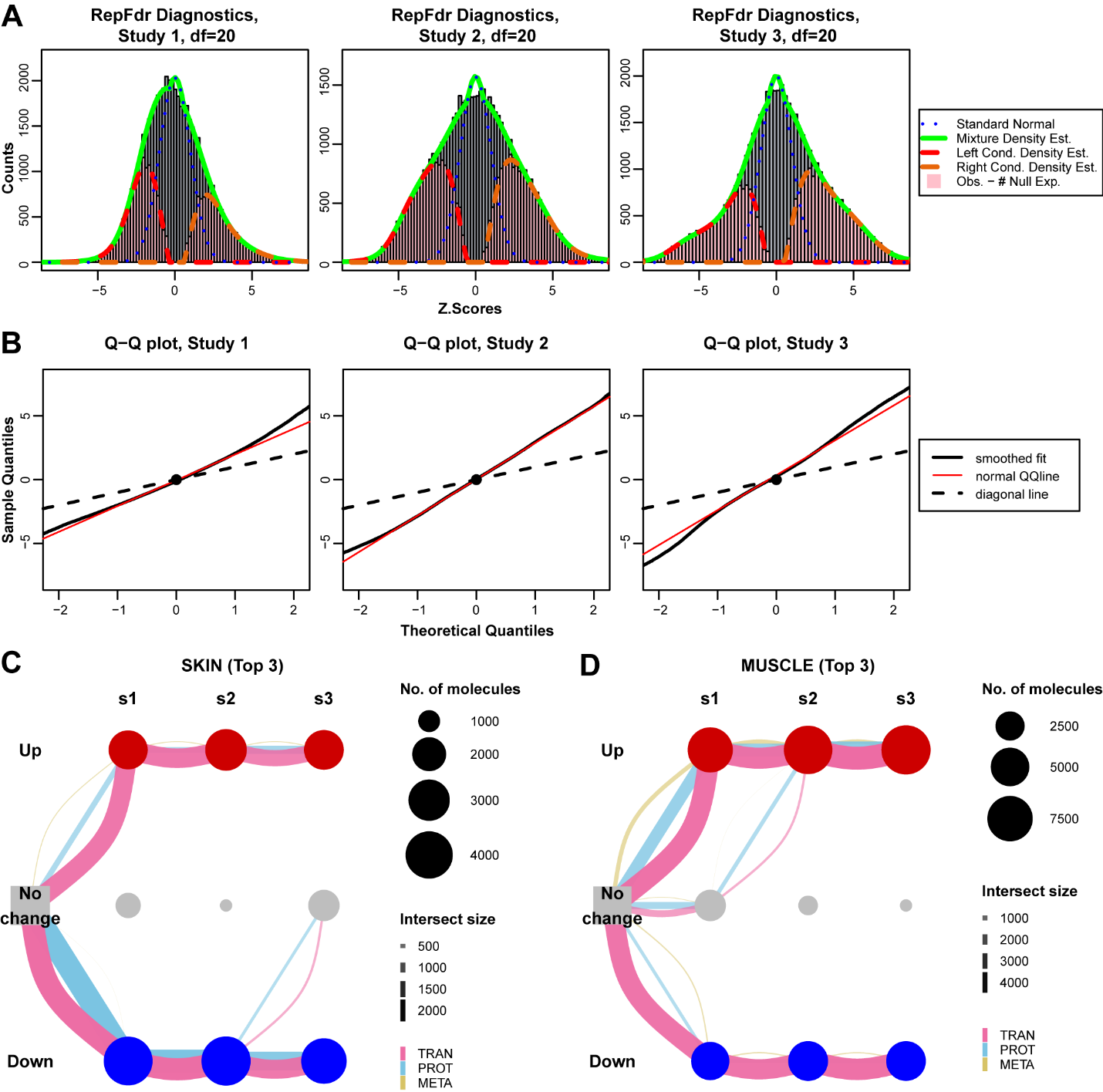

**Fig. S9. Repfdr diagnostic plots and visualization of the top three trajectories.** (**A**) Histogram of observed z-scores (grey bars) with multiple density estimation curves: blue dashed line represents standard normal density, green solid line denotes fitted overall mixture density estimate, red and orange dashed lines represent left-tail and right-tail conditional density estimates, respectively; pink shading highlights deviation of observed distribution from global null hypothesis. (**B**) Quantile-quantile (QQ) plot comparing empirical z quantiles with theoretical standard normal quantiles; black dashed line represents the theoretical diagonal (perfect fit to standard normal), red solid line is the normal QQ reference curve, and black solid line is the smoothing curve for empirical quantiles, used to assess tail deviation and model fit quality. Model parameters: df = 20; n.bins = 150; central.prop = 0.25. (**C**) Top three major temporal trajectories of skin injury-differentiated molecules (visualizing trajectory flow and molecular count per stage). (**D**) Top three major temporal trajectories of muscle injury-differentiated molecules. Node size in **C** and **D** indicates the number of molecules in that stage (column), while edges represent molecular trajectories over time, with edge width proportional to the number of molecules in that trajectory.

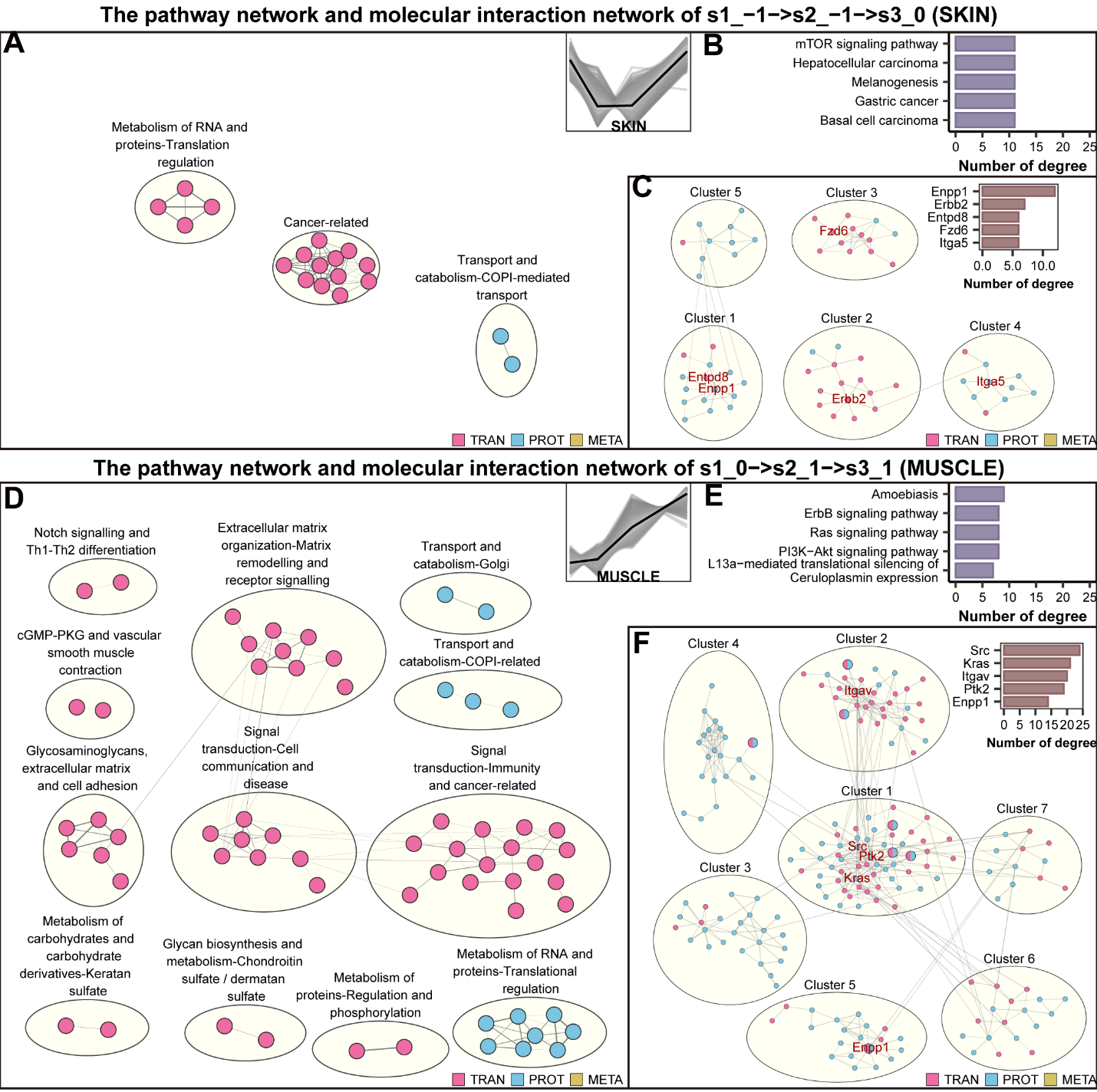

**Fig. S10. Pathway networks and molecular interactions for the third trajectory of skin and muscle.** (**A**) Skin third trajectory (SKIN: s1_-1 -> s2_-1 -> s3_0): pathway network of significantly enriched pathways. Each node represents a pathway; node color indicates the omics category to which it belongs; node size is proportional to the number of omics categories involved in the pathway. Edges represent pairs of functionally similar pathways (similarity score ≥ 0.5), with edge thickness proportional to the similarity between nodes. Pathway nodes were clustered using the Louvain algorithm to define biological themes. (**B**) Top five pathway nodes by degree in panel **A**. (**C**) Molecular interaction subnetwork for SKIN third-trajectory features derived from a prior-knowledge interaction network. Each node represents a molecule, with node color indicating its omics category and node size proportional to the number of omics categories the molecule belongs to. Edges denote highly correlated molecular pairs (similarity score ≥ 0.5), with edge thickness proportional to the similarity between nodes. Modules identified by leading-eigenvector clustering. (**D-F)** Corresponding panels for the MUSCLE third trajectory (MUSCLE: s1_0 -> s2_1 -> s3_1); annotations and conventions are as in **A-C**.

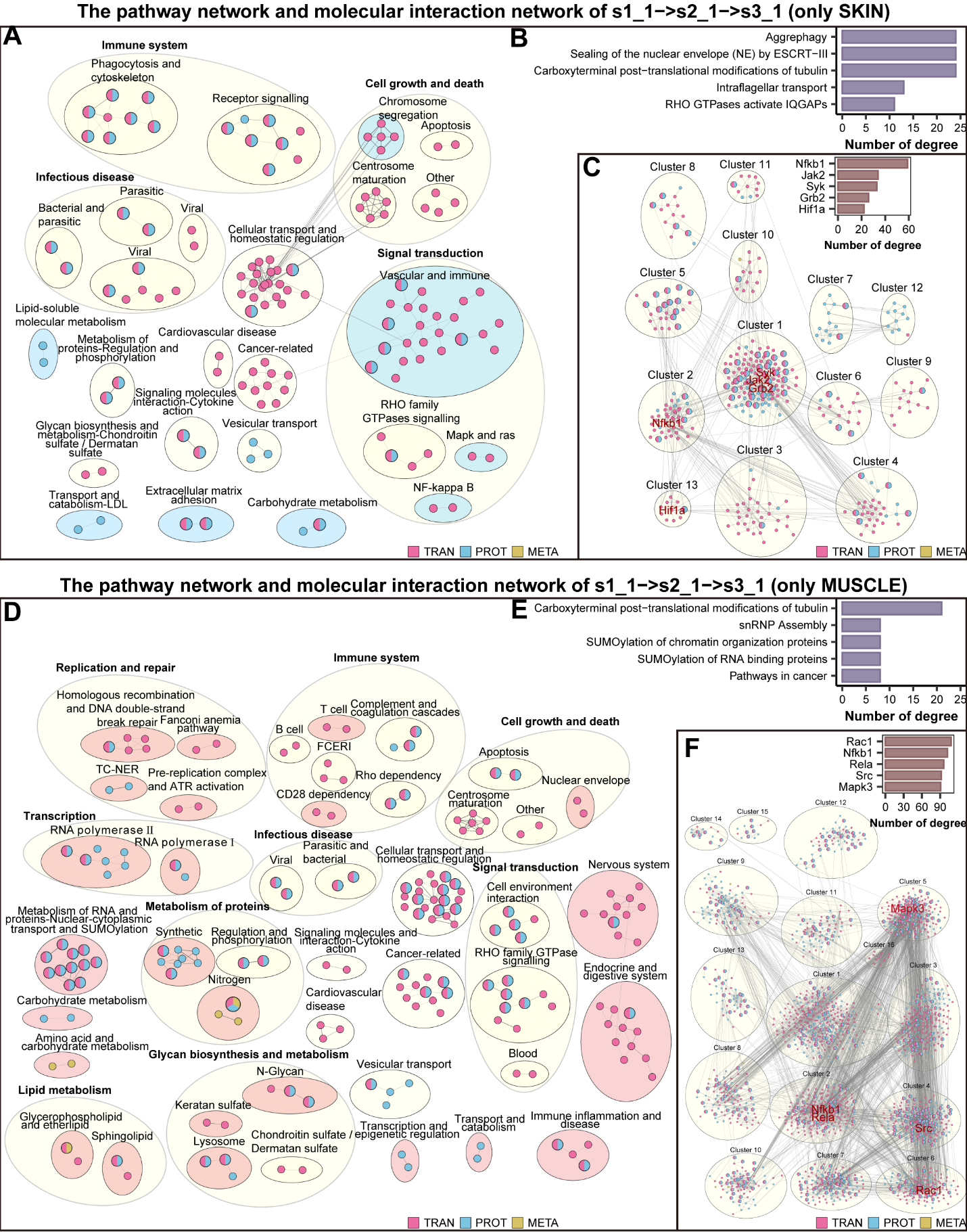

**Fig. S11. Pathway networks and molecular interactions of skin and muscle sustained up-regulation trajectories.** (**A**) Skin sustained up-regulated trajectory (s1_1 -> s2_1 -> s3_1): pathway network (node, edge conventions as **Fig. S10A**). Light-blue shading marks skin-specific pathways in this network. (**B**) Top five high-degree pathway nodes in panel **A**. (**C**) Molecular interaction subnetwork for skin sustained up-regulated features (node, edge conventions as **Fig. S10C**). (**D-F)** Analogous panels for muscle sustained up-regulated trajectory (s1_1 -> s2_1 -> s3_1); light-red shading denotes muscle-specific pathways.

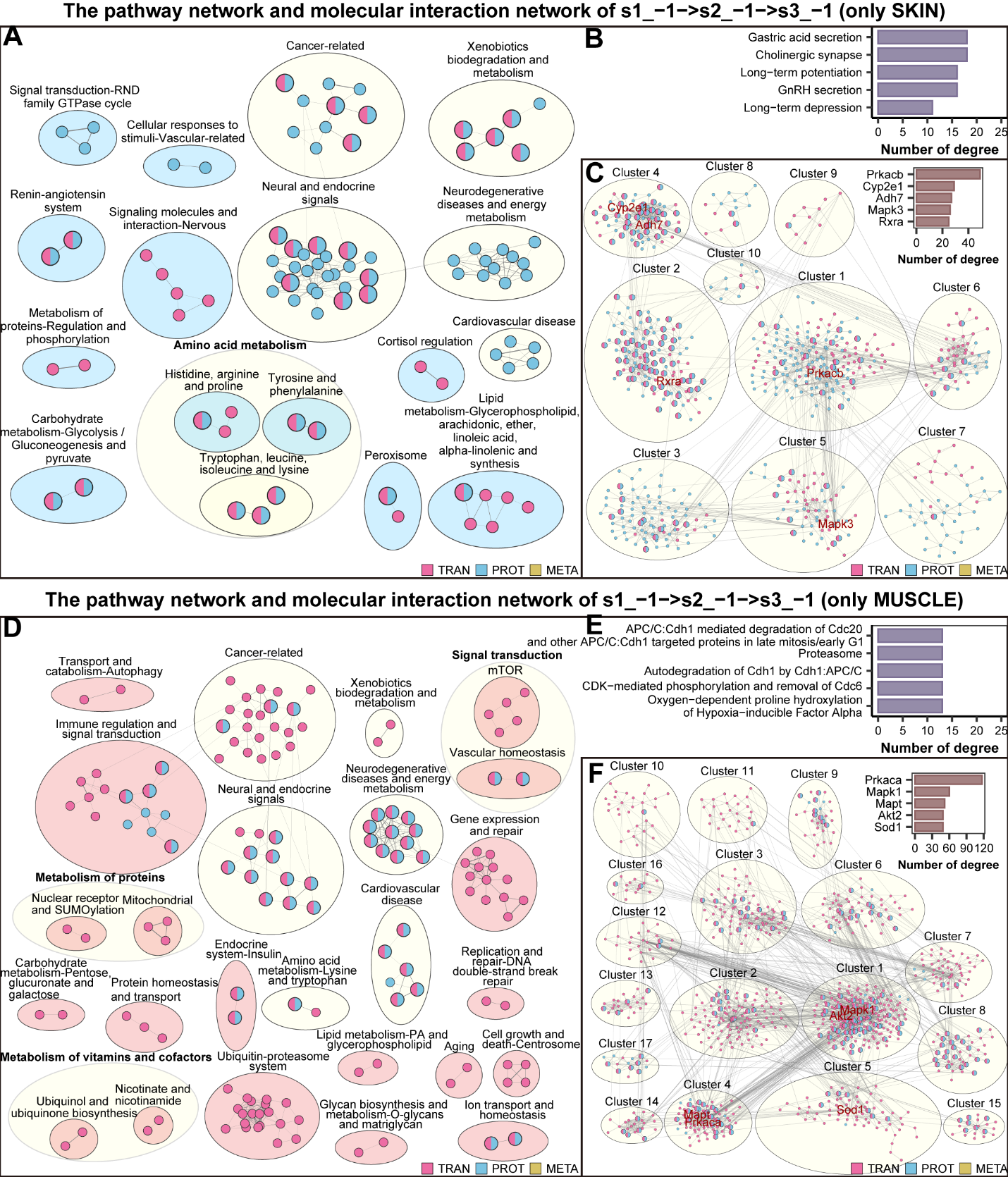

**Fig. S12. Pathway networks and molecular interactions for sustained down-regulation trajectories in skin and muscle.** (**A**) Skin sustained down-regulated trajectory (s1_−1 -> s2_−1 -> s3_−1): pathway network (node, edge conventions as **Fig. S10A**). Light-blue shading marks skin-specific pathways. (**B**) Top five high-degree pathway nodes in panel **A**. (**C**) Molecular interaction subnetwork for skin sustained down-regulated features (node, edge conventions as **Fig. S10C**). (**D–F)** Corresponding panels for muscle sustained down-regulated trajectory (s1_−1 -> s2_−1 -> s3_−1); light-red shading denotes muscle-specific pathways.

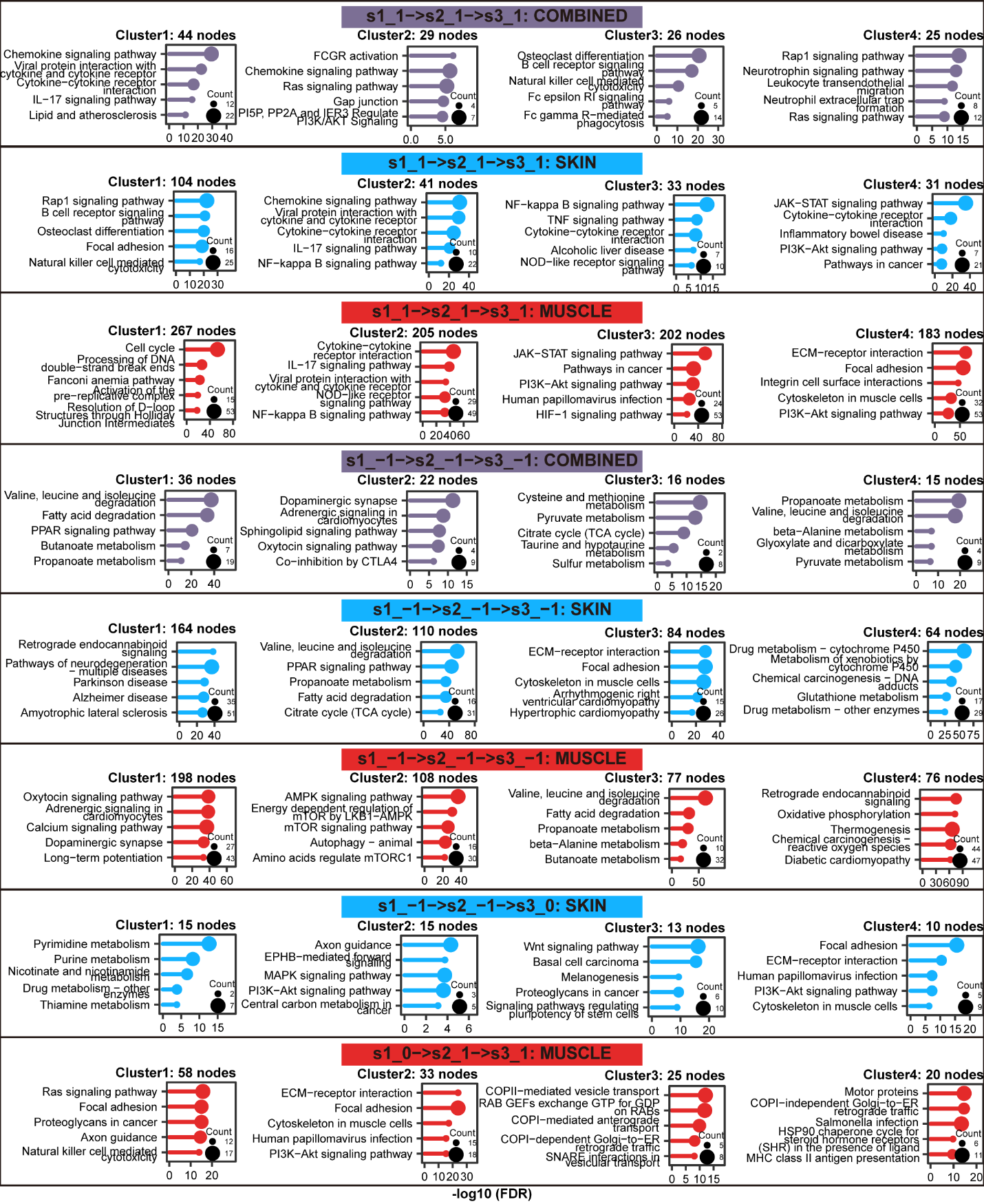

**Fig. S13. Pathway enrichment analysis for molecular interaction network clusters across different trajectories (FDR < 0.05).** From top to bottom: (1) skin and muscle sustained up-regulated shared subnetwork; (2) skin sustained up-regulated subnetwork; (3) muscle sustained up-regulated subnetwork; (4) skin and muscle sustained down-regulated shared subnetwork; (5) skin sustained down-regulated subnetwork; (6) muscle sustained down-regulated subnetwork; (7) skin third-trajectory subnetwork; (8) muscle third-trajectory subnetwork. Each row displays pathway enrichment results for the four largest clusters (based on FDR < 0.05 significance threshold) after partitioning the corresponding subnetwork into modules (clustering clusters). Each subgraph is annotated with the names of the top five significantly enriched pathways, their enrichment significance, and the number of corresponding molecules.

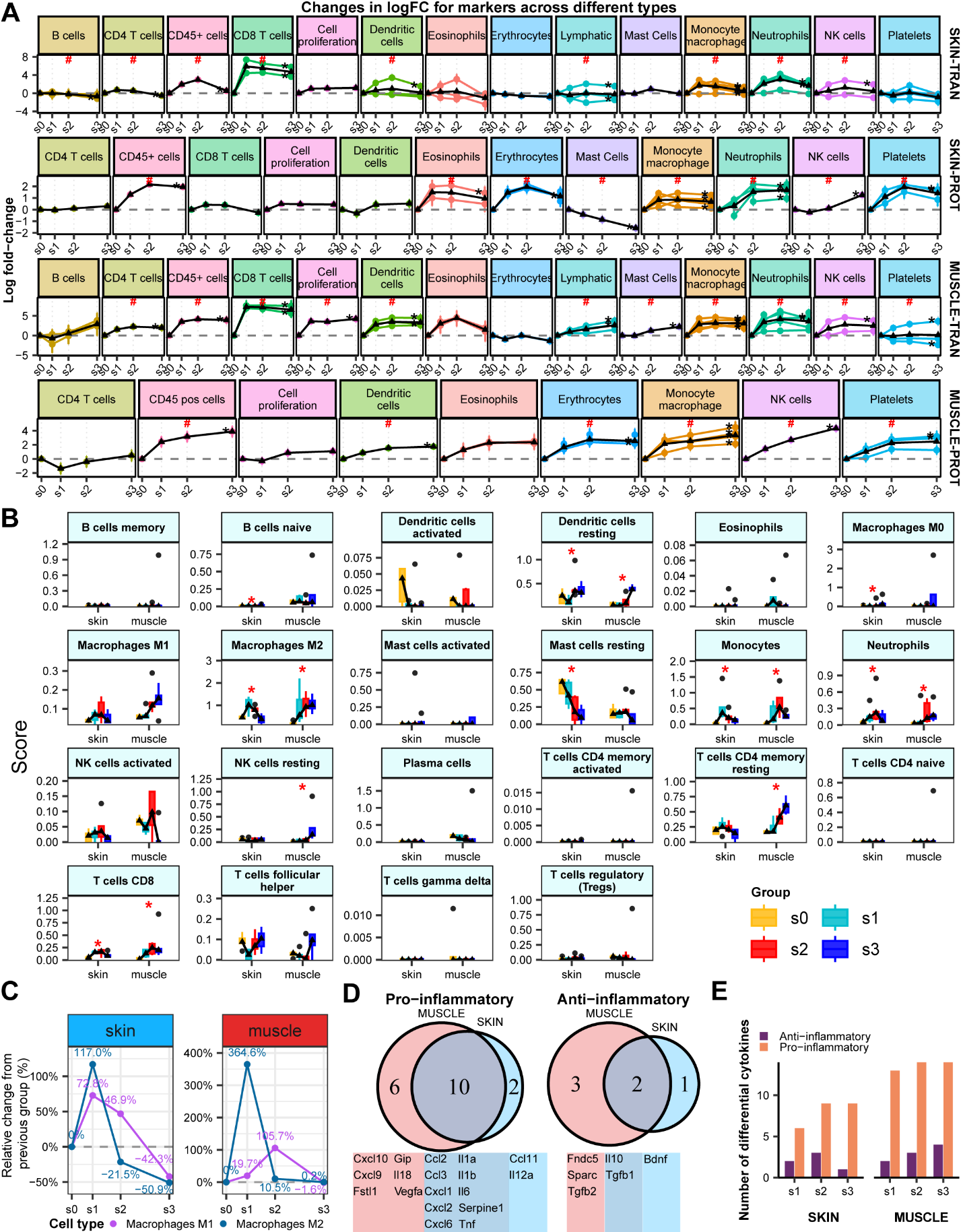

**Fig. S14. Effects of pressure injury on immune responses.** (**A**) Log2FC changes (transcript and protein) for immune-cell marker genes in skin and muscle across stages. Data points in the line chart represent the log2FC of each molecule at the corresponding stage (log2FC at s0 set to 0), with error bars indicating the standard error of log2FC (lfcSE); the black line denotes the average log2FC for the indicated molecule set. Symbol # indicates at least one feature in the set is significant (FDR < 0.05). (**B**) Boxplots of absolute scores for 22 immune cell types inferred by CIBERSORTx. Box plots follow standard definitions (Q1, median, Q3; upper/lower bars represent max/min; filled points indicate outliers). The line chart displays the median absolute score for each cell type across groups. Multi-group comparisons use Kruskal-Wallis test; significance threshold two-sided p < 0.05 (*p < 0.05). (**C**) Stagewise relative growth rate of M1 and M2 macrophage absolute scores (relative to the previous stage). (**D**) Venn diagram of pro-inflammatory and anti-inflammatory factors that change significantly (FDR < 0.05) during the injury process; the study used 18 pro-inflammatory and 6 anti-inflammatory factors. (**E**) Bar chart showing counts of significantly differential pro- and anti-inflammatory factors per tissue and stage (FDR < 0.05).

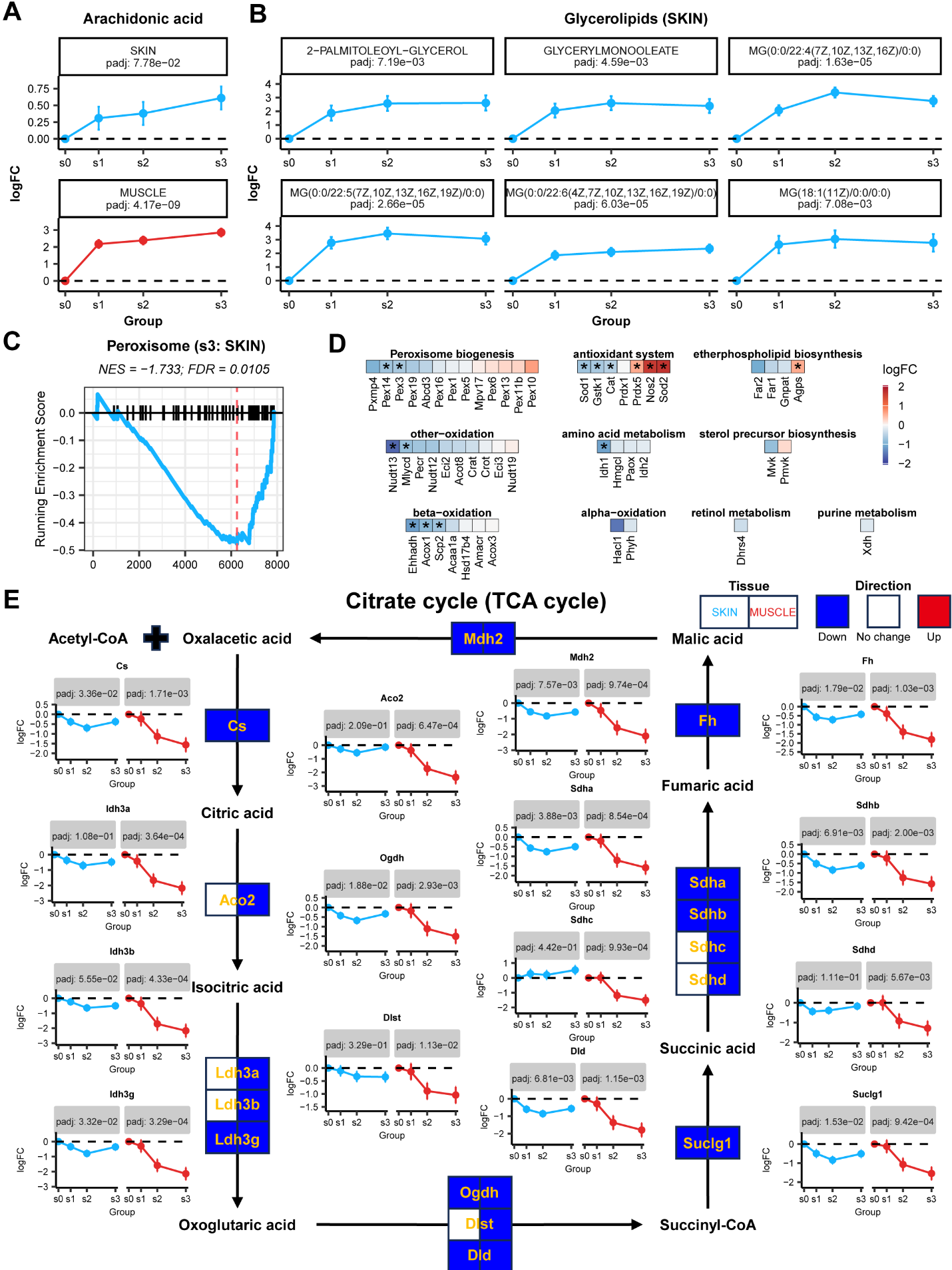

**Fig. S15. Effects of pressure injury on metabolic responses.** (**A**) Log2FC changes of arachidonic acid (AA) across tissues and stages. (**B**) Log2FC changes of six significantly altered (FDR < 0.05) glyceride molecules in skin. (**C**) GSEA result for peroxisome pathway at skin s3. (**D**) Protein log2FC changes for detected members of the peroxisome pathway at skin s3. *FDR < 0.05. (**E**) Changes of selected TCA pathway proteins in skin (left) and muscle (right). Box color coding: red indicates significantly upregulated proteins (FDR < 0.05 and log2FC > 0); blue indicates significantly downregulated proteins (FDR < 0.05 and log2FC < 0); white indicates no significant change. Line chart annotations as in **Fig. S14A**.

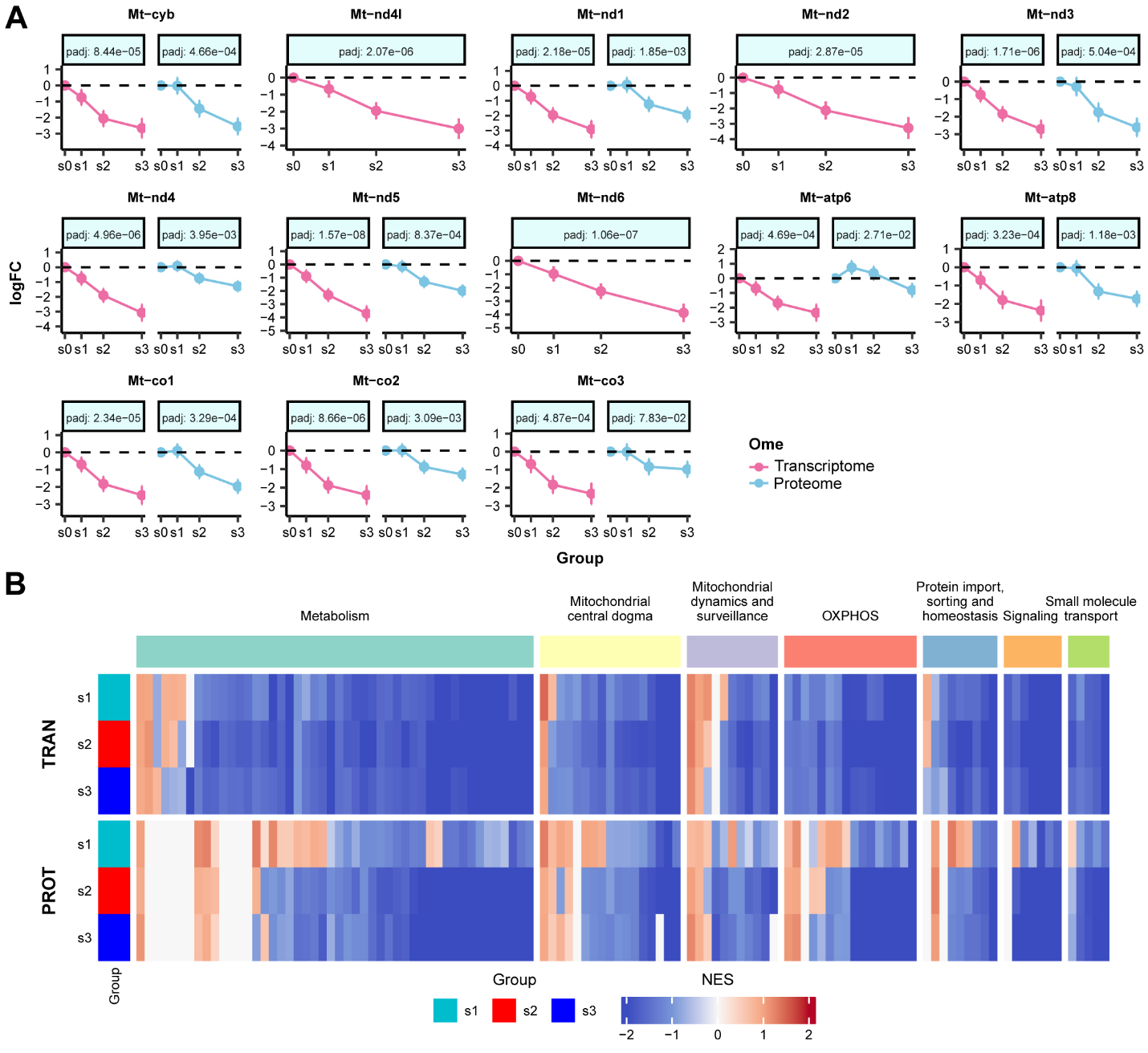

**Fig. S16. Effects of pressure injury on muscle mitochondrial-related responses.** (**A**) Log2FC trajectories across stages for 13 mitochondrially encoded genes at transcript and protein levels. Points represent log2FC values (with s0 set to 0), and error bars denote lfcSE. (**B**) Heatmap of GSEA normalized enrichment scores (NES) for MitoCarta-defined mitochondrial pathways across stages and layers (TRAN: transcriptome; PROT: proteome); pathways are grouped by MitoPathways classification.

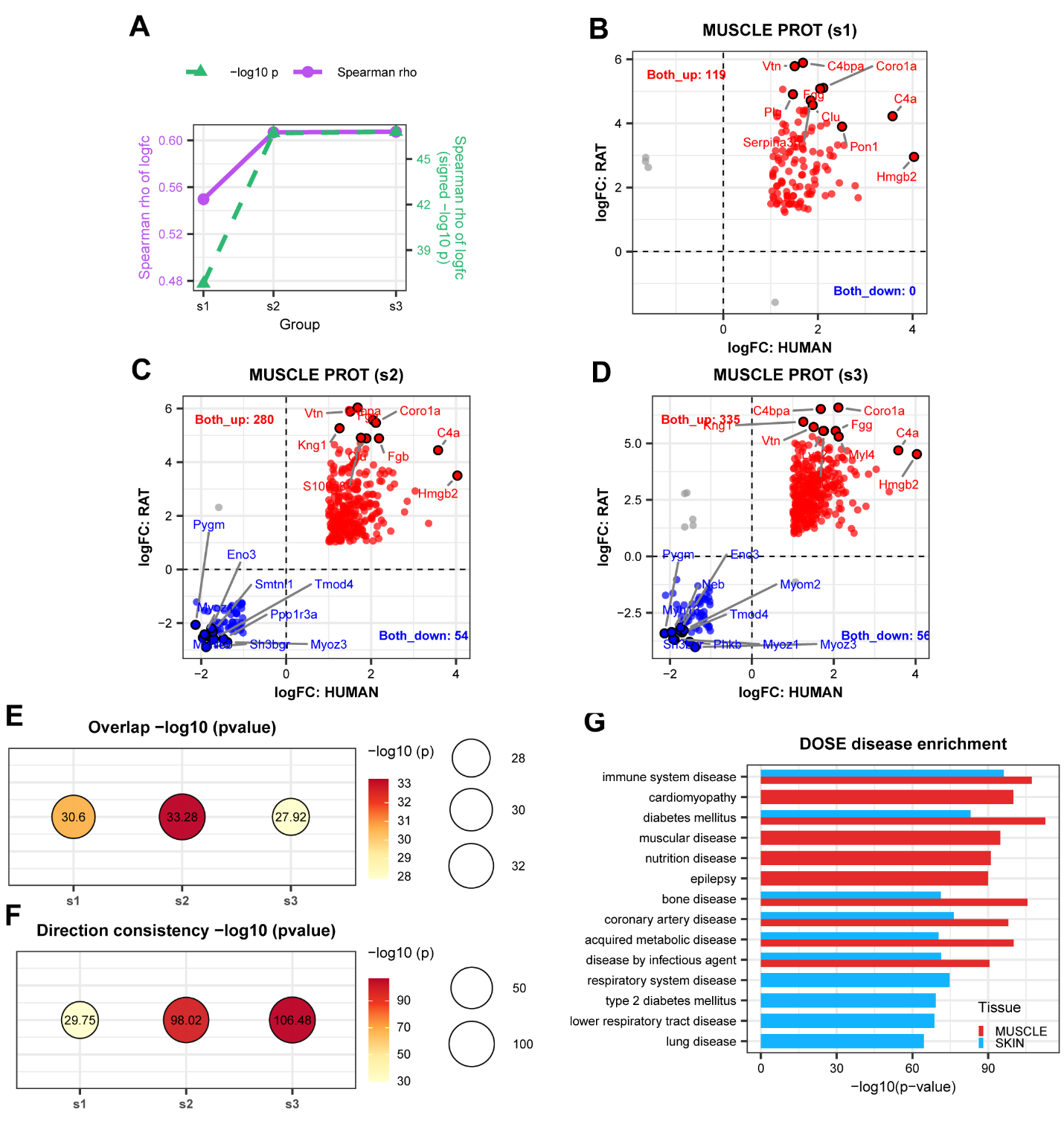

**Fig. S17. Comparison and similarity analysis with human pressure injury data.** (**A**) Spearman correlation between rat differential proteins and Liu et al. human PI differential proteins (log2FC). (**B**–**D**) Scatter plots of log2FC distribution for overlapping differential proteins between rat (stages s1, s2, s3) and human comparisons: red dots indicate proteins upregulated in both species; blue dots indicate proteins downregulated in both species; gray dots indicate proteins with inconsistent directionality. The top ten feature proteins with the largest average log2FC upregulation or downregulation across both species are highlighted. (**E**) Fisher’s exact test for significance of overlap between species’ differential proteins. (**F**) Exact binomial test for directional concordance among overlapping differential proteins. (**G**) Disease ontology (DO) enrichment for skin and muscle differential features: top ten significantly enriched disease terms (FDR < 0.05).
